# A relational architecture of the human genome

**DOI:** 10.64898/2026.10.15.759350

**Authors:** Jeremias Krause, Martin Danner, Florian Kraft, Tanhim Islam, Matthias Begemann, Zain Ziad, Miriam Elbracht, Mathis Bode, Ingo Kurth

## Abstract

Complex systems exhibit organization not only through their components but also via their interdependencies. Local nucleotide dependencies underpin regulatory motifs and RNA structures, yet principles governing a higher-order organization across the human genome remain unknown. Here we systematically perturbed every position of the complete telomere-to-telomere human genome using Nucleotide Transformer v3 with an exascale compute cluster, generating nucleotide-resolution dependency maps that reveal a directed, weighted network of position-to-position genomic relationships. This network recapitulates known biological features and uncovers recurrent, directional architectures termed ‘relons’, spanning coding, non-coding, and unannotated regions. These relational modules reflect evolutionary and functional constraints and associate with human genetic variation and pathogenic variants. Our findings suggest a higher-order relational layer of genome organization, providing a framework to interpret DNA as a complex system of dependencies beyond linear and 3D structure, with implications for understanding genetic variation and disease.

---

Complex systems are found throughout nature and society, including cities, economies, the Internet, ecosystems, immune systems, or the human brain. Despite their diverse components and functions, these systems share common organizational principles. One such principle is nonlinear behaviour, where the effect of perturbing one component depends on the state of others^1–3^. For example, identical carbon atoms can form fundamentally different materials depending on their arrangement and connectivity, as seen in graphite and diamond. Similarly, waves such as sound or light can amplify or cancel each other depending on their relative phase. Interactions among many elements can generate emergent properties and self-organized structures that are not apparent from the components in isolation^1,2,4–6^. Many complex systems are also adaptive, with variation and selection reshaping their organization over time^5^. Thus, understanding these systems requires considering both their individual components and the relationships that connect them, since local dependencies generate patterns and larger structures that define the operational context and interrelations among components^1–3,5–7^.

The human genome exemplifies a complex system, with its organization resolved across multiple dimensions. The telomere-to-telomere (T2T) reference provides an essentially complete linear sequence map, while pangenome efforts increasingly capture human genomic diversity^8,9^. Functional genomics and projects such as ENCODE have mapped regulatory and epigenetic states across cellular contexts^10^. Three-dimensional genomics has revealed physical relationships between distant loci through compartments, topologically associating domains (TADs), and chromatin loops^11–13^, and sequence-based models increasingly predict aspects of this organization directly from DNA^14,15^. Additionally, genetic-interaction and co-essentiality networks reveal functional relationships between genes, organizing them into complexes, pathways and cellular processes^16,17^. Collectively, these approaches have transformed our view of the genome from a linear sequence into a multilayered biological system. However, at the DNA level, we still primarily describe positions and elements individually, rather than the architecture of dependencies between them. Genomic language models have been shown to learn contextual properties of DNA sequence^18–28^, while systematic in silico perturbations of sequence-based neural networks have revealed cis-regulatory interactions^29^. Local nucleotide dependencies are also well established, for example, bases within transcription-factor binding motifs can affect binding interdependently, and genomic language models have recently been shown to recover dependencies underlying regulatory motifs and coordinated splice signals^30,31^. However, it remains unclear whether such dependencies are isolated features or if higher-order dependencies constitute a genome-wide organizational system. This knowledge gap raises the question of whether a relational layer of organization exists at the nucleotide level across the entire genome.

Here, we use Nucleotide Transformer v3^32^ and large-scale GPU-computing to systematically perturb the complete T2T human reference genome *in silico*. We substitute each of approximately 3.12 billion positions with each of the three alternative nucleotides and record the resulting changes in model predictions at other positions within the surrounding sequence context, capturing position-to-position responses. This approach enables us to construct nucleotide-resolution dependency maps and represent the human genome as a genome-scale weighted relational graph. Our analysis indicates that these local relationships can be represented as a higher-order dependency architecture spanning the entire genome. To facilitate exploration of this relational organization, we provide a genome-wide relational representation through an openly accessible genome browser (https://relome.org/). Together, our results suggest a relational layer of genome organization and establish a framework for understanding DNA as a complex system of dependencies, with important implications for interpreting human variation and disease-causing variants.

## Results

### Genome-wide dependency maps

Genome organization is increasingly resolved at the level of individual sequence elements and three-dimensional chromatin contacts, yet genomic positions are still largely interpreted independently at the sequence level. A complementary perspective is to ask how the state of one genomic position relates to that of another and if a resulting network captures biologically meaningful organization (**Fig. 1**). Such dependencies can be quantified using genomic language models by perturbing individual nucleotides and measuring the resulting changes in model predictions at other positions. Recent work by da Silva *et al*.^30^ formalized nucleotide dependency analysis and demonstrated that dependency maps recover regulatory motifs, RNA structural contacts, and other functional sequence relationships, establishing their utility as an interpretable readout of genomic language models. Here, we asked whether nucleotide dependencies, when mapped across the complete human genome, form a higher-order architecture of their own. To address this, we generated nucleotide dependency maps across the complete T2T human reference genome (T2T-CHM13v2.0). The 3,117,275,501-bp genome was partitioned into 8-kb sequence windows, yielding 761,065 windows. These windows were offset by 4 kb to ensure overlapping sequence contexts and minimize boundary effects. Within each window, every nucleotide position was evaluated in its reference state and after substitution with each of the three alternative nucleotides (perturbation). We used the Nucleotide Transformer v3 (NTv3) pre-model, a transformer-based DNA language model trained on genomic sequences to learn context-dependent patterns and relationships between nucleotides. We quantified how each perturbation altered model responses at all other positions within the same 8-kb sequence context, generating a nucleotide-resolution dependency map for every analysed window. In total, this required 24,938,643,456 model forward passes, performed on the exascale supercomputer JUPITER^33^ using approximately 150,000 GPU hours on NVIDIA H100 GPUs. Each dependency map was represented as a directed, weighted graph in which genomic positions are *nodes* and detected dependencies are *edges*. The direction of each edge specifies the perturbed source position and the responding target position, while the edge weight reflects dependency strength. Notably, these relationships are directional: perturbing nucleotide position A and measuring the response at nucleotide position B does not necessarily yield the same dependency as perturbing B and measuring the response at position A. Our representation captured both the strength and direction of relationships between positions. Collectively, these maps transformed the genome from a collection of individually annotated positions into a genome-scale network of nucleotide relationships, providing a framework for analysing connectivity, directionality, and higher-order genomic sequence organization. To reduce their visual complexity and reveal interpretable relational patterns, we transformed the dependency maps into arc diagrams, made accessible through the openly accessible Relome browser (https://relome.org/) (**Fig. 1**).

**Fig. 1.**
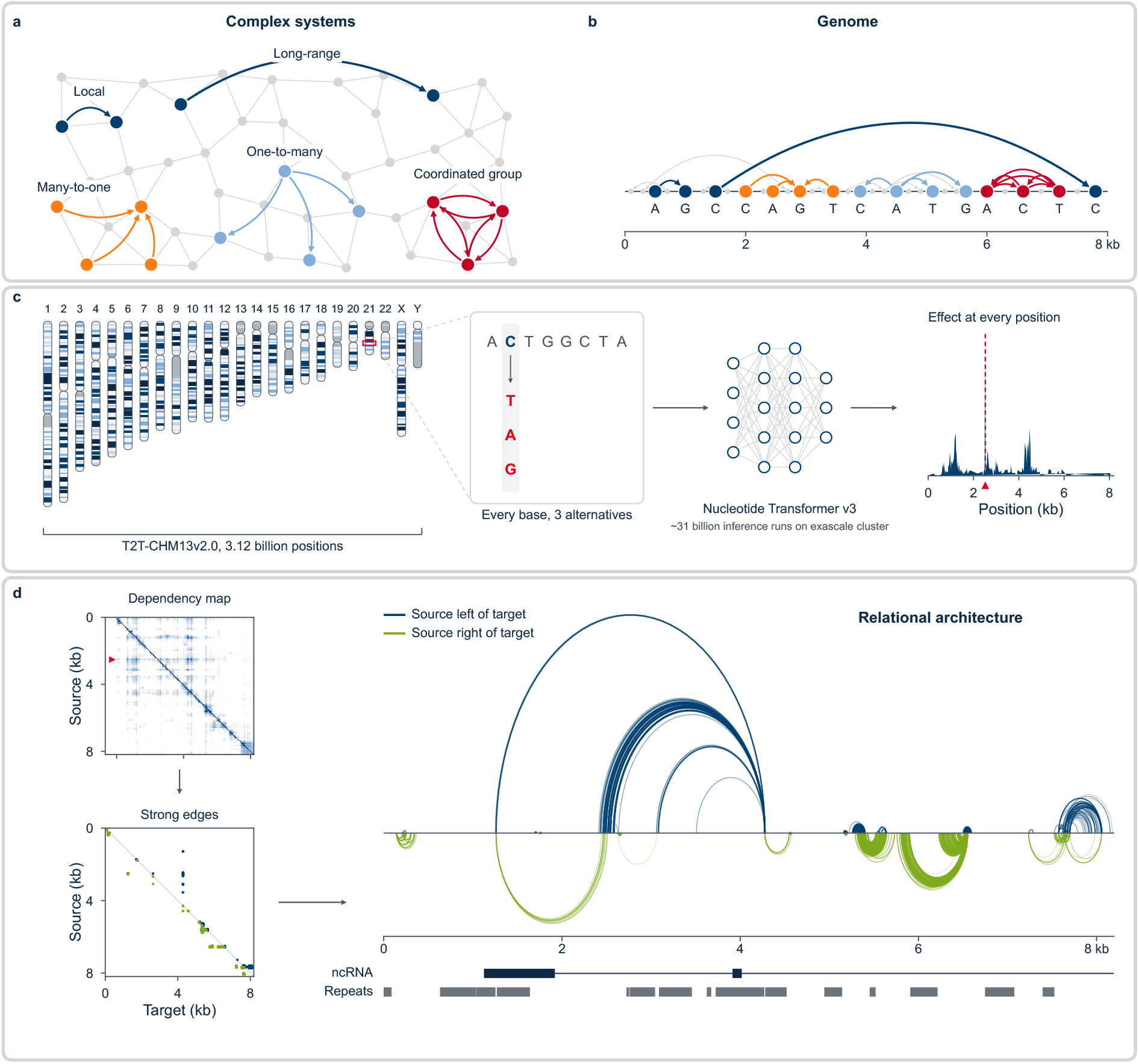
Genome-wide nucleotide dependency mapping. **a,** Schematic representation of complex systems as networks of interacting components, characterized by a nonlinear, high-dimensional architecture in which the effect of one component depends on the state of others, giving rise to local and long-range dependencies, many-to-one and one-to-many relationships, and coordinated groups of interconnected elements. **b,** Conceptual representation of the human genome as a complex system, in which dependencies between nucleotides may follow general organizational principles similar to those observed in other complex systems. **c,** Genome-wide *in silico* mutagenesis of the T2T-CHM13v2.0 human reference genome (approx. 3.12 billion positions). Each nucleotide was systematically substituted with all three alternative bases, and Nucleotide Transformer v3 was used with large-scale GPU computing to quantify the resulting changes in model predictions at other positions within 8-kb sequence windows. **d,** Transformation of nucleotide-dependency matrices into interpretable genomic representations. Strong dependencies were extracted from directed, weighted matrices (source positions on the vertical axis, target positions on the horizontal axis) and visualized as arcs along the genomic sequence. Blue arcs indicate dependencies from lower to higher genomic coordinates, green arcs the reverse; opacity reflects dependency strength. Genomic annotations, including non-coding RNAs and repeats, provide biological context.

### Dependencies reflect genomic features

We first asked whether nucleotide dependencies reflect established biological properties of genomic sequence. Using phyloP^34^ ≥2.7 to define evolutionarily conserved positions, we found that dependencies preferentially connected conserved nucleotides across all genomic distances (**Fig. 2a-c**). To account for the clustering of conserved positions and the strong distance dependence of nucleotide relationships, enrichment was assessed against candidate position pairs matched for genomic distance within 500-kb blocks. Estimates are from fixed-effect meta-analysis across the 22 autosomes. For edge counts, conserved-to-conserved dependencies were enriched 1.91-fold (95% CI, 1.88– 1.94) at 1–99 bp, increasing to 23.3-fold (22.4–24.1) at 4–5 kb and remaining 10.1-fold (9.2–11.1) at 8.0–8.2 kb. Dependencies involving only one conserved position showed intermediate enrichment, whereas those between non-conserved positions remained close to expectation. Dependency weights followed a similar pattern (**Fig. 2c**).

**Fig. 2.**
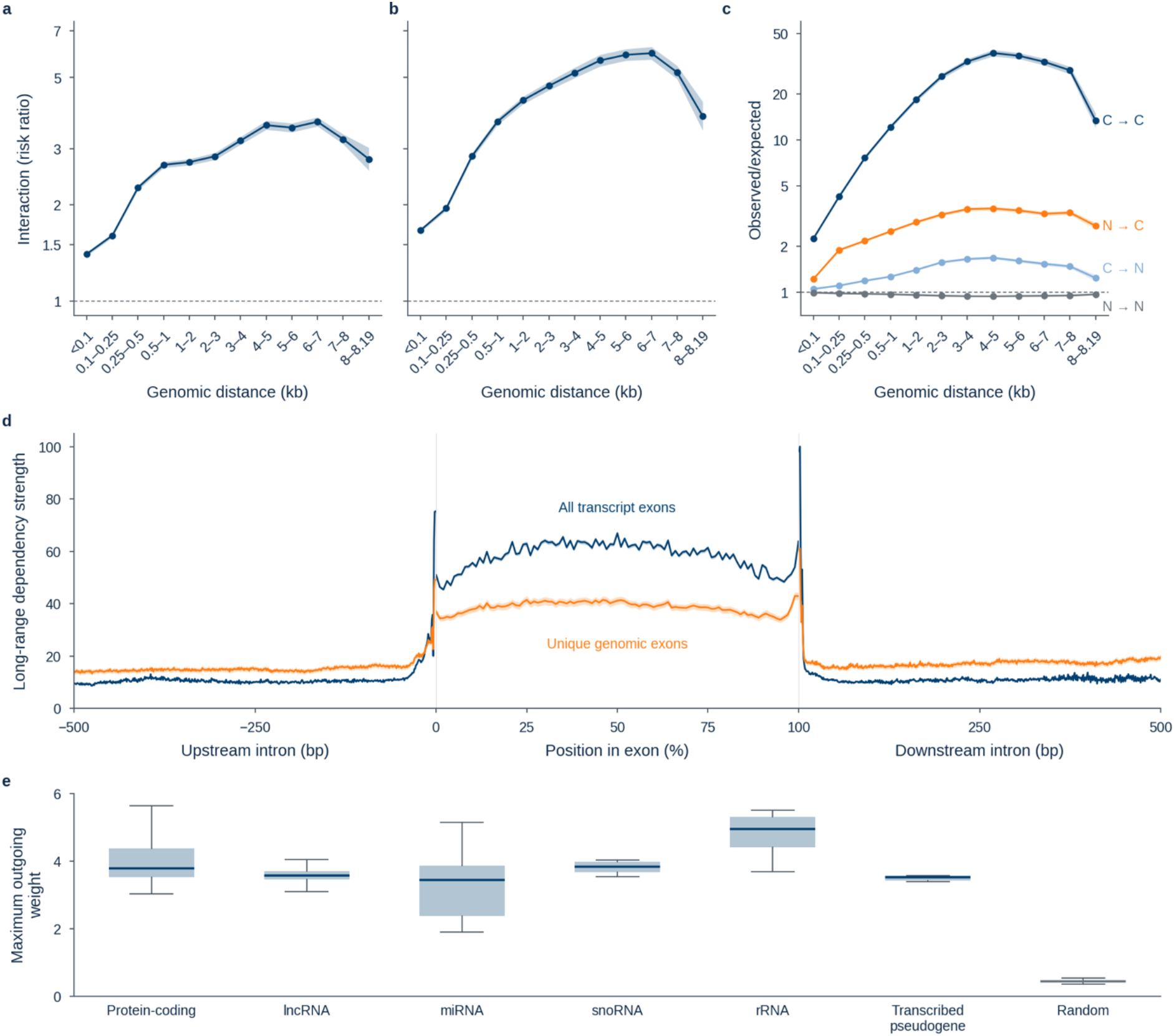
Nucleotide dependencies reflect genomic features. **a,b**, Conservation-specific interaction across genomic distances for edge counts (a) and edge weights (b), measuring preferential conserved-to-conserved coupling beyond the independent effects of source and target conservation. **c**, Observed-to-expected enrichment of dependency weights for conserved (C; phyloP ≥2.7) and non-conserved (N) source–target pairs. Expected values account for genomic distance and candidate-pair composition within 500-kb blocks. **a–c**, Lines show pooled estimates across autosomes; shaded regions indicate 95% confidence intervals. Dashed lines indicate no enrichment or interaction. **d**, Mean long-range dependency strength (≥2 kb) across chromosome 21 exons, scaled to relative exon length (0–100%), and 500-bp flanking introns. All transcript exons and unique genomic exons are shown separately; vertical lines indicate exon boundaries. **e**, Distribution of maximum outgoing dependency weights across protein-coding genes, non-coding RNA classes and transcribed pseudogenes, compared with 140 synthetically generated random 8-kb sequences. For each nucleotide position, the maximum weight corresponds to its strongest outgoing dependency.

Importantly, conserved nucleotides preferentially connected to one another beyond their individual propensity to form dependencies. This conservation-specific interaction increased from 1.40-fold (95% CI, 1.39–1.42) at 1–99 bp to 3.55-fold (3.44–3.67) at 4–5 kb and 3.64-fold (3.52–3.76) at 6–7 kb (**Fig. 2a**). Dependency weights showed an even stronger interaction, reaching 5.96-fold (5.69–6.24) at 6–7 kb (**Fig. 2b**). Beyond 2 kb, these effects were consistent across autosomes and robust to alternative phyloP thresholds (**Supplementary Fig. 2**). Thus, evolutionary constraint is reflected not only in individual nucleotides but also in the organization and strength of their relationships. Furthermore, the observed effects were reproducible, across different phyloP thresholds (**Supplementary Fig. 1**).

We next examined whether dependencies also capture gene structure. On chromosome 21, long-range dependency strength (≥2 kb) increased sharply at exon boundaries and remained elevated throughout exons relative to flanking introns, both across all transcript exons and after merging overlapping exons into unique genomic intervals (**Fig. 2d**). Finally, we compared the strongest outgoing dependency at each nucleotide position across annotated genomic elements and 140 synthetically generated random 8-kb sequences. Maximum dependency weights were substantially higher in biological sequences than in random controls, with the highest values observed in rRNA (**Fig. 2e**). Together, these findings demonstrate that nucleotide dependencies reflect evolutionary constraint and characteristic features of genomic organization, providing a biological foundation for investigating their higher-order architecture.

### Dependencies follow distinct rules

We next asked whether nucleotide dependencies follow reproducible organizational rules across the genome. After filtering low-confidence edges (Methods), we examined their variation with genomic separation, chromosomal context and source–target annotation. Dependency persistence differed markedly between genomic compartments (**Fig. 3a**). In gene-rich and gene-poor regions, retention declined steeply with distance, reaching 1.8% and 1.0%, respectively, at 4–8 kb. By contrast, 41% and 60% of possible pairs were retained in heterochromatic and centromeric regions. Median edge weight declined approximately 45-fold with distance, whereas the strongest dependencies persisted over kilobase scales (**Fig. 3b**). Much of the elevated long-range retention in heterochromatic and centromeric regions reflected repetitive sequences. After excluding repeat-associated pairs, retention at 4–8 kb fell to 2.0%, approaching the 1.3% of repeat-free pairs in gene-rich regions. Nevertheless, weights of the strongest 0.1% of dependencies increased from 1.4 at 257–512 bp to 3.4 at 4–8 kb (**Supplementary Figure 3**). We therefore defined strong edges as the top 0.1% within genomic separation and chromosomal context strata. Strong dependencies connected genomic elements non-randomly (**Fig. 3c,d**). Coding sequence was preferentially targeted at short distances across gene-rich and gene-poor regions, whereas several promoter-associated pairings were depleted. These associations were context dependent, with some pooled enrichments driven by individual loci rather than recurrent chromosome-wide effects. Thus, dependency architecture reflects genomic distance, chromosomal environment and sequence identity rather than a uniform interaction rule. These global trends and locus-specific patterns prompted us to investigate whether dependencies form recurrent higher-order structures.

**Fig. 3.**
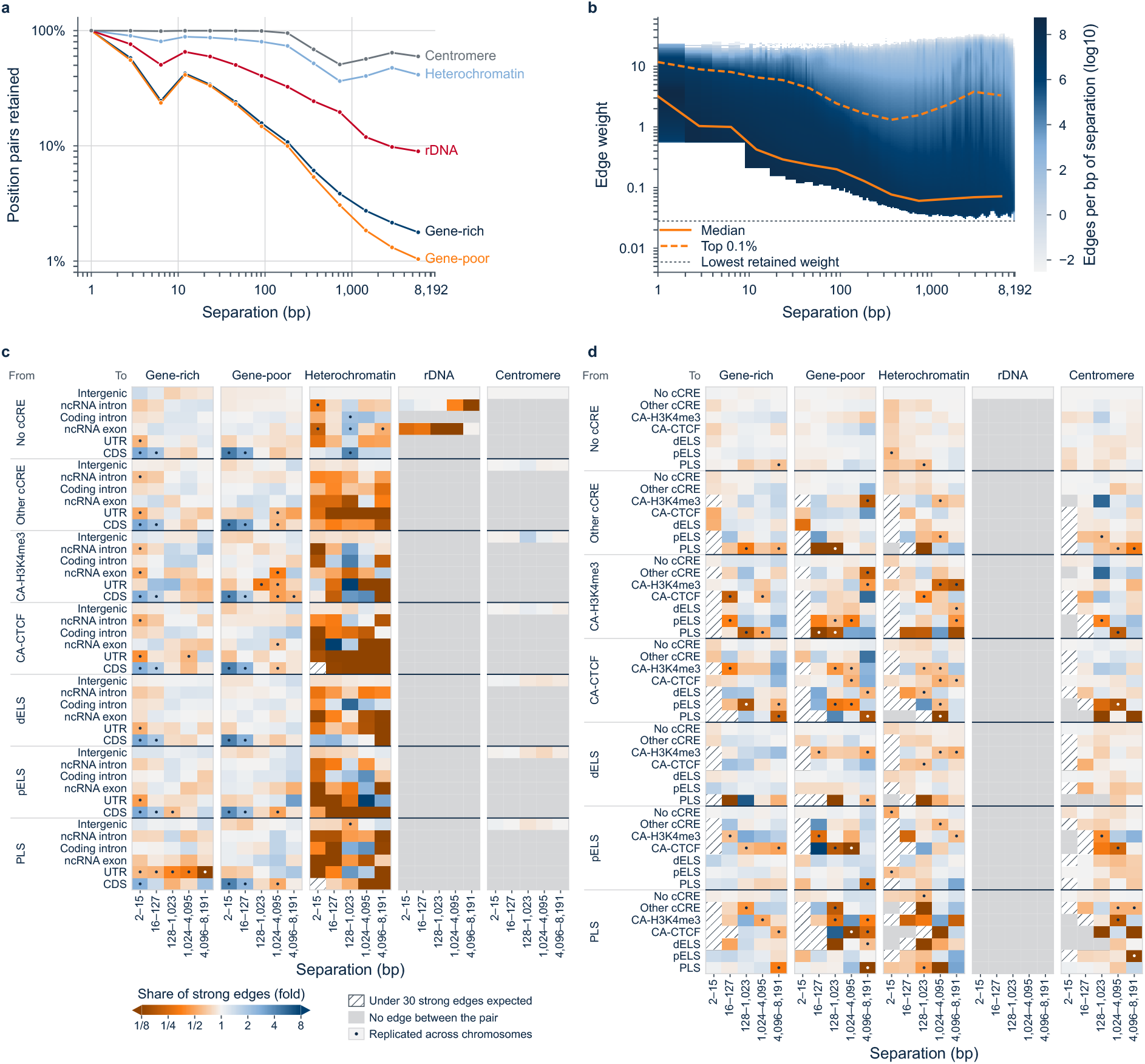
Nucleotide dependencies follow distinct organizational rules. **a,** Fraction of position pairs retaining a dependency after filtering, stratified by genomic separation and cytoband class. **b,** Distribution of retained edge weights with increasing genomic separation; lines indicate the median and top 0.1% of edges. **c,d,** Enrichment or depletion of strong edges between cCRE and gene-element classes (**c**) and between cCRE classes (**d**), relative to dependencies matched for separation and cytoband context. Strong edges represent the top 0.1% of retained edges within each separation and cytoband stratum. Dots indicate effects of at least 1.5-fold that were directionally consistent across chromosomes; hatched cells indicate limited expected counts and grey cells absence of eligible edges.

### Recurrent relational architectures

We next asked whether nucleotide dependencies organize into recurrent structural units across the genome. Analysis of the strongest 0.1% of dependencies revealed distinct, non-random patterns of nucleotide connectivity. We systematically identified regions enriched for strong dependencies relative to a background accounting for genomic distance and cytoband class (**Fig. 4a**). Across the genome, these regions resolved into seven characteristic geometries, which we term Relons (for *relational organization*). Relons are defined by their dependency architecture rather than nucleotide sequence, and similar structures can be detected in unrelated sequences. The seven classes capture distinct forms of nucleotide organization (**Fig. 4b**). *Blocks* represent densely interconnected stretches, whereas *periodic* Relons exhibit dependencies at recurring nucleotide intervals. Three classes describe relationships between two sequence stretches: *offset diagonals* connect positions in the same order, *anti-diagonals* connect them in reverse order, and *islands* link stretches without a consistent positional correspondence. Because dependencies are directional, perturbing one nucleotide need not affect another to the same extent as the reverse perturbation. Accordingly, *source hubs* and *target hubs* describe asymmetric relationships in which a narrow sequence stretch influences, or depends on, a substantially broader region. Together, these seven geometries capture local connectivity, periodicity, long-range correspondence and directional dependencies. Each produces a recognizable pattern in the Relome browser (**Fig. 4c**).

**Fig. 4.**
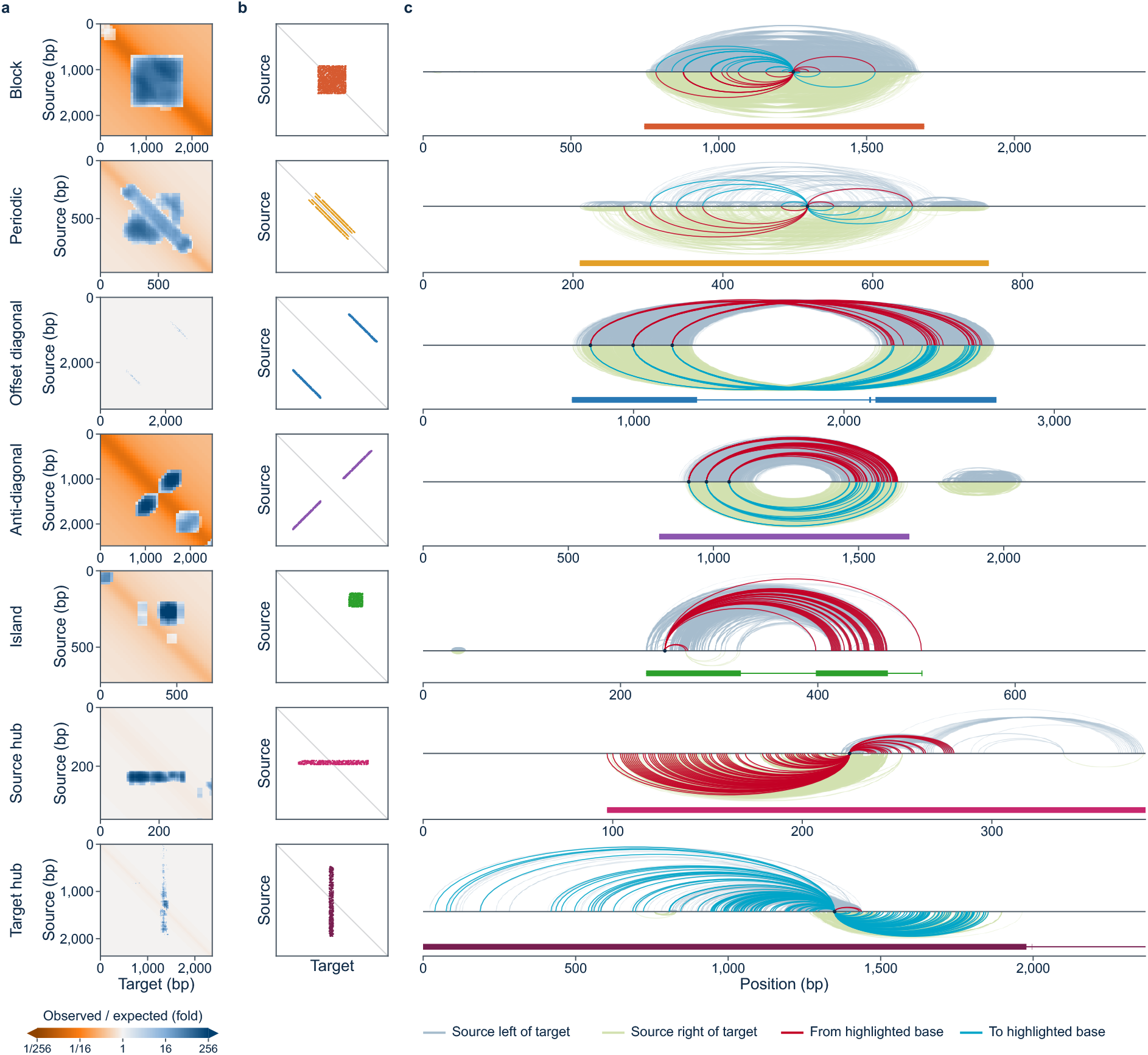
Detection and classification of recurrent Relon architectures. Representative examples of the seven Relon classes from chromosome 21 (T2T-CHM13), with one class per row. **a,** Detection of enriched dependency structures in 8,192-bp nucleotide-dependency maps. Source (perturbed) and target (responding) nucleotides are shown on the vertical and horizontal axes, respectively. Heatmaps show the log₂ observed-to-expected ratio of strong edges within square boxes of the indicated size, calculated as log₂[(observed + 0.5)/(expected + 0.5)]. Expected edge counts account for nucleotide separation and cytoband class; strong edges comprise the top 0.1% within these strata. Blue indicates enrichment and green depletion relative to expectation. **b,** Idealized dependency geometries defining the seven classes. Blocks form dense diagonal squares with edges spanning multiple distances; periodic Relons concentrate at least 50% of their edges at the three most frequent separations. Offset diagonals connect separate sequence stretches in the same positional order (correlation ≥ 0.5), whereas anti-diagonals exhibit reversed correspondence (correlation ≤ −0.5). Islands connect separate stretches without positional correspondence. Source hubs connect a narrow source interval to a target interval at least four times wider; target hubs exhibit the reverse organization. Dependencies are directional, and reciprocal edges are assessed independently. Islands and hubs are illustrated in one direction; the remaining classes in both. **c,** Corresponding regions in the Relome browser (https://relome.org/). Blue arcs above the sequence axis indicate dependencies towards higher genomic coordinates, whereas green arcs below indicate dependencies towards lower coordinates. Arc opacity reflects dependency strength; red and cyan highlight outgoing and incoming edges, respectively, for the selected nucleotide. The underlying track indicates the detected Relon and its class.

### Relons form a genome-wide landscape

We next examined how Relons are distributed across the human genome and whether their organization follows common patterns across chromosomes. We compared two complementary edge filters: the strongest 0.1% of dependencies and the strongest 1% after excluding edges involving repetitive sequences or non-B DNA motifs (**Fig. 5**; Methods). Non-B DNA refers to alternative DNA conformations that alter the local physical structure of the canonical double helix^35^. These identified 1,286,832 and 1,208,023 Relons, respectively, with all seven classes present on every chromosome. Relon distribution varied markedly between genomic regions (**Fig. 5a**). Under the 0.1% filter (strongest 0.1% of dependency edges), Relons covered a median of 12% of chromosome bases (range 8–60%), with the highest coverage on chromosome Y. Centromeric and heterochromatic regions were dominated by relatively few, large Relons, covering 95% and 73% of their sequence, respectively. Euchromatin, by contrast, contained approximately 440 discrete Relons per Mb, covering 7.6% of its bases. Coverage was largely uniform along chromosome arms but increased to 13.5% within the terminal 2 Mb. Under the second filter (strongest 1% of remaining edges after excluding dependencies with either endpoint in a RepeatMasker^36^ element or non-B DNA motif), Relons covered a median of 21% of unmasked chromosome bases, while euchromatic Relon density remained similar (416 per Mb) and the increase towards chromosome ends persisted (25.1% versus 20.4%). Despite these regional differences, Relon class composition was remarkably consistent across chromosomes (**Fig. 5b**). Under the 0.1% filter, anti-diagonals were most frequent (33.5%), followed by islands (17.2%), blocks (15.8%), periodic Relons (11.6%), target hubs (10.8%), source hubs (6.4%) and offset diagonals (4.8%). No class varied by more than two percentage points in standard deviation across chromosomes. The second filter substantially shifted these proportions, particularly reducing anti-diagonals to 14.9% and increasing blocks to 32.6%, yet class composition remained similarly consistent across autosomes. Because the filters differed in both sequence exclusion and edge threshold, these shifts cannot be attributed to repetitive sequences alone. Thus, Relons exhibit regional differences in size and coverage alongside relatively stable class proportions across chromosomes.

**Fig. 5.**
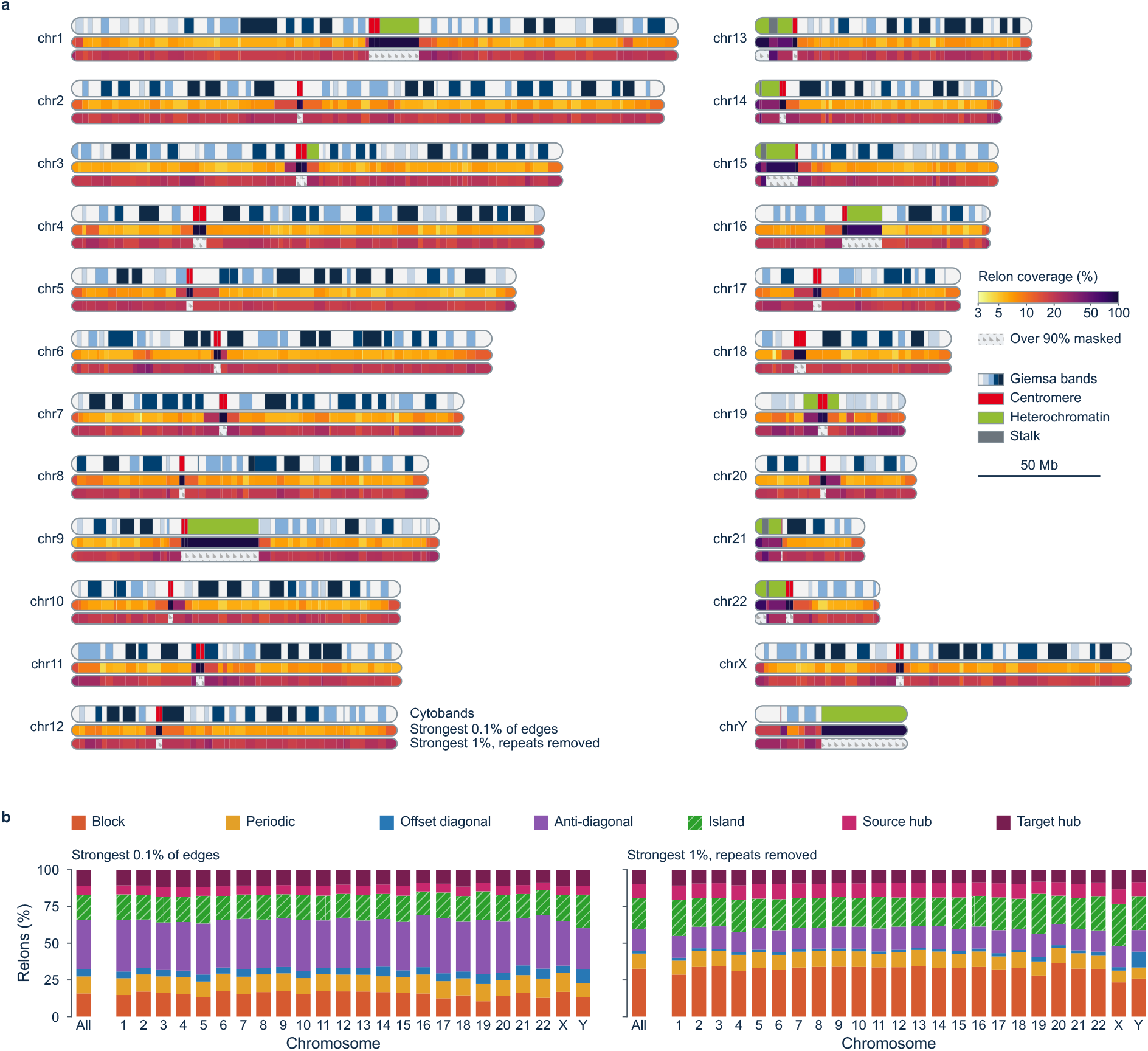
Chromosomal distribution and class composition of Relons. Genome-wide distribution of Relons across all 24 chromosomes of the T2T-CHM13 reference genome, assessed using two edge filters. **a,** Relon coverage per cyto-band, defined as the fraction of bases contained within Relon stretches and displayed beneath chromosome ideograms. The upper track shows Relons identified using the strongest 0.1% of dependency edges, selected separately by nucleotide separation and cytoband class. The lower track shows Relons identified using the strongest 1% of remaining edges after excluding dependencies with either endpoint in a RepeatMasker element or non-B DNA motif, retaining RNA genes and their repeat-annotated copies. For the latter filter, coverage is calculated relative to unmasked bases; cytobands with more than 90% masked sequence are indicated by dots. **b,** Relative abundance of the seven Relon classes across individual chromosomes and genome-wide under both edge filters. Genome-wide class proportions are indicated above each panel.

### Relons within and beyond annotation

We next asked whether Relon architectures reflect known genomic features and whether they also occur beyond current annotation. To avoid dominance by large centromeric and heterochromatic Relons, enrichment analyses were restricted to euchromatin (**Fig. 6a**). Under the strongest 0.1% edge filter, most Relon classes showed limited enrichment for RepeatMasker elements (0.9–1.5-fold). In contrast, non-B DNA motifs, covering 9.4% of euchromatin, accounted for 62% of periodic Relon bases (6.6-fold enrichment) and 35% of anti-diagonal bases (3.8-fold). After excluding repeat-associated and non-B DNA dependencies and selecting the strongest 1% of remaining edges, coding sequence was enriched in blocks, periodic Relons and islands (4.3–4.9-fold). These enrichments persisted when individual Relon candidates were analysed before merging overlapping structures (**Supplementary Fig. 4**). Notably, source hubs also showed strong coding-sequence enrichment at the candidate level (8.9-fold), indicating that its absence among merged source hubs resulted from the merging procedure rather than a lack of association with coding sequence.

**Fig. 6.**
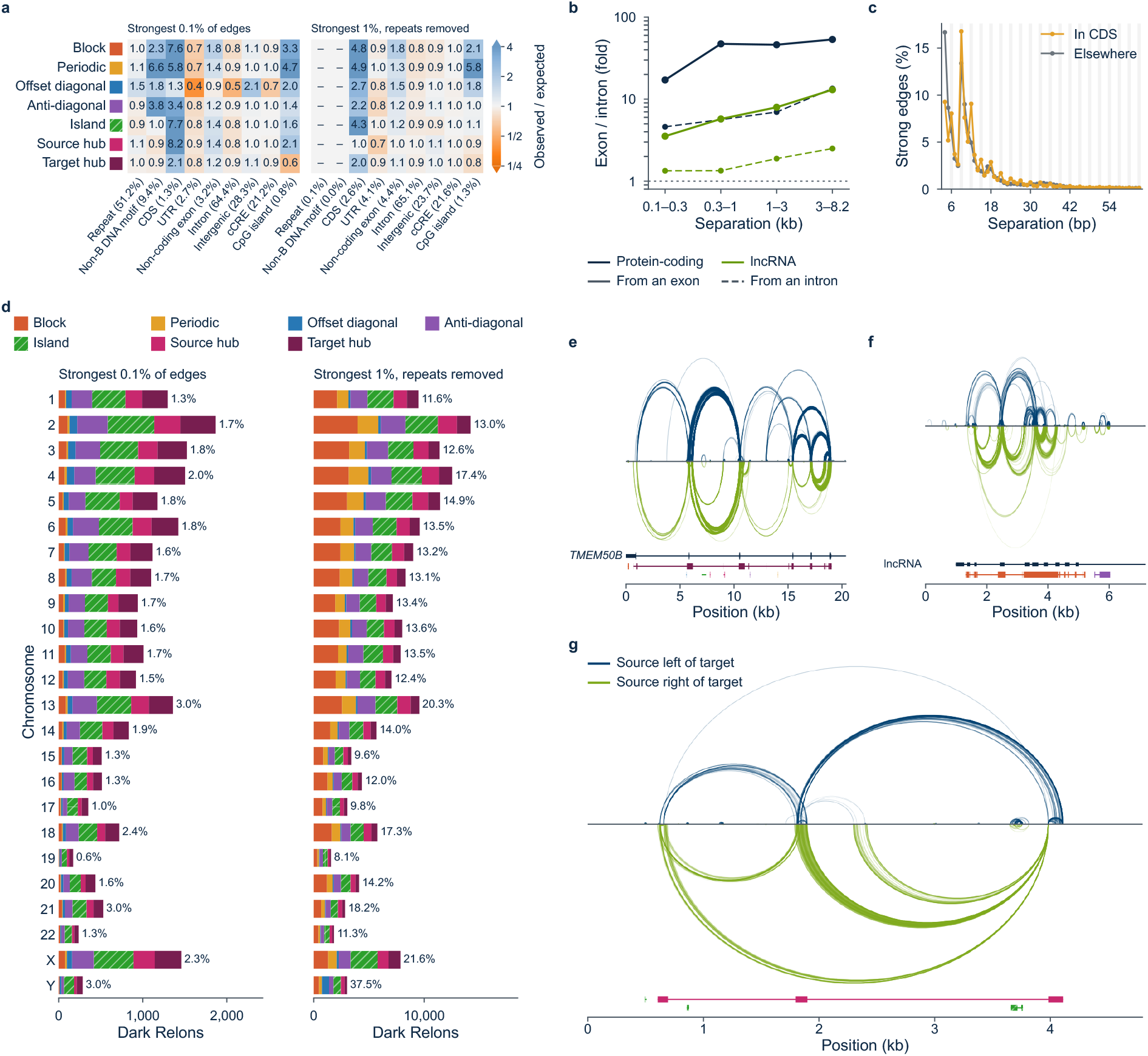
Relon architectures within and beyond genomic annotation. **a,** Enrichment of genomic elements across Relon classes, expressed as observed-to-expected ratios of overlapping bases relative to euchromatic sequence. Heatmaps show the strongest 0.1% of dependency edges (left) and the strongest 1% after excluding repeat-associated and non-B DNA dependencies (right). Colours indicate log₂ enrichment (blue, depleted; red, enriched). **b,** Exon-to-exon dependency enrichment relative to exon-to-intron dependencies at matched genomic distance (solid lines), with corresponding intron-to-exon versus intron-to-intron ratios as controls (dashed lines). **c,** Nucleotide separations (4–63 bp) of strong edges in periodic Relon candidates predominantly overlapping coding sequence (≥50% of bases) versus other candidates, repeat-excluded filter. Multiples of three are highlighted; n indicates candidate counts. **d,** Dark Relons, defined by the absence of overlapping genomic annotations, shown by chromosome and class under both edge filters. Percentages indicate their proportion among all Relons per chromosome; panels have separate count scales. **e-f**, Protein-coding and long non-coding RNA genes are shown separately, using the repeat-excluded filter and edges within the same Relon. **g,** Representative Dark Relon [R27721, source hub, chr9:95,115,985–95,119,490, strong 1% no-repeat] in the Relome browser, showing its dependency architecture alongside genomic annotation tracks. Blue and green arcs indicate dependencies towards higher and lower genomic coordinates, respectively; opacity reflects edge weight. All analyses use T2T-CHM13; panels a–d cover all 24 chromosomes. Additional methodological details and class-specific analyses are provided in the Supplementary Results.

We next examined whether Relons also capture gene structure (**Fig. 6b,c,e,f**). At matched genomic distances beyond 300 bp, exon-to-exon dependencies were 46–53 times denser than exon-to-intron dependencies in protein-coding transcripts, with corresponding enrichments of 3.5–13-fold across distance ranges in long non-coding RNA genes (**Fig. 6b**). These preferences exceeded analogous intron-based controls by 4–8-fold and approximately 3–5-fold, respectively. Relon dependencies also reflected the coding reading frame: 47.6% of edges in periodic candidates predominantly overlapping coding sequence occurred at separations divisible by three, compared with 33.8% in other candidates (**Fig. 6c**). Both exon connectivity and three-base periodicity were more pronounced under the unmasked 0.1% edge filter, with 54.7% versus 34.4% of edges occurring at three-base intervals (**Supplementary Fig. 5**). Thus, without using gene annotations for their detection, Relons recover characteristic features of gene organization, including exon connectivity and the three-base periodicity of coding sequence.

Finally, we asked whether recurrent architectures also occur in regions without known genomic elements. We term these Dark Relons, defined by the absence of overlap with annotated genes, repeats, candidate regulatory elements, small open reading frames, non-B DNA motifs, tandem repeats or CpG islands. Under the strongest 0.1% filter, we identified 22,220 Dark Relons (1.7% of all Relons), spanning every chromosome and all seven classes, with islands and source or target hubs accounting for 68% (**Fig. 6d**). Under the repeat-excluding 1% filter, 169,103 Relons (14.0%) were dark, partly reflecting the greater proportion of unannotated sequence remaining after masking. To assess whether Relons preferentially occur in unannotated regions, we compared them with copies randomly shifted within the same cytoband class. Under the 0.1% filter, Dark Relons were less frequent than expected (1.7% versus 4.9%), consistent with the concentration of Relons in annotated repeats and motifs. After repeat exclusion, however, this depletion disappeared: among Relons without masked bases, 17.6% were dark, compared with 17.0% of their shifted controls. Thus, although Relons are not generally enriched in unannotated sequence, tens of thousands of recurrent dependency architectures occur beyond existing genome annotation. A representative source hub (R27721, chr9:95,115,985– 95,119,490) illustrates such a structure (**Fig. 6g**). These findings extend the relational landscape beyond known genomic elements, identifying previously uncharacterized dependency structures for further investigation.

### Dependencies reflect human variation

Having established that Relons capture known genomic organization and extend beyond current annotation, we next asked whether nucleotide dependencies are associated with human genetic variation and disease. We first examined clinically classified variants in ClinVar^37^. Positions harbouring pathogenic variants showed substantially more outgoing (2.15-fold, 95% CI, 2.09–2.22) and incoming dependencies (2.73-fold, 2.65–2.81) than benign positions (**Fig. 7a**). This difference persisted after accounting for genomic context and evolutionary conservation, including among phyloP-adjusted missense positions (outgoing, 1.52-fold, 95% CI, 1.40–1.66; incoming, 1.48-fold, 1.36–1.61; **Fig. 7b**). Although pathogenic positions also preferentially connected to one another, this largely reflected their greater overall connectivity. After accounting for genomic distance and the independent effects of source and target pathogenicity, their mutual edge counts were close to expectation (median observed-to-expected ratio, 1.04; range, 0.98–1.08). Their dependency weights, however, remained 1.19-fold higher than expected (95% CI, 1.17– 1.22) across all 22 autosomes (**Supplementary Fig. 7**). Thus, pathogenic positions are characterized by increased connectivity and stronger dependencies, rather than a distinct disease-specific network.

**Fig. 7.**
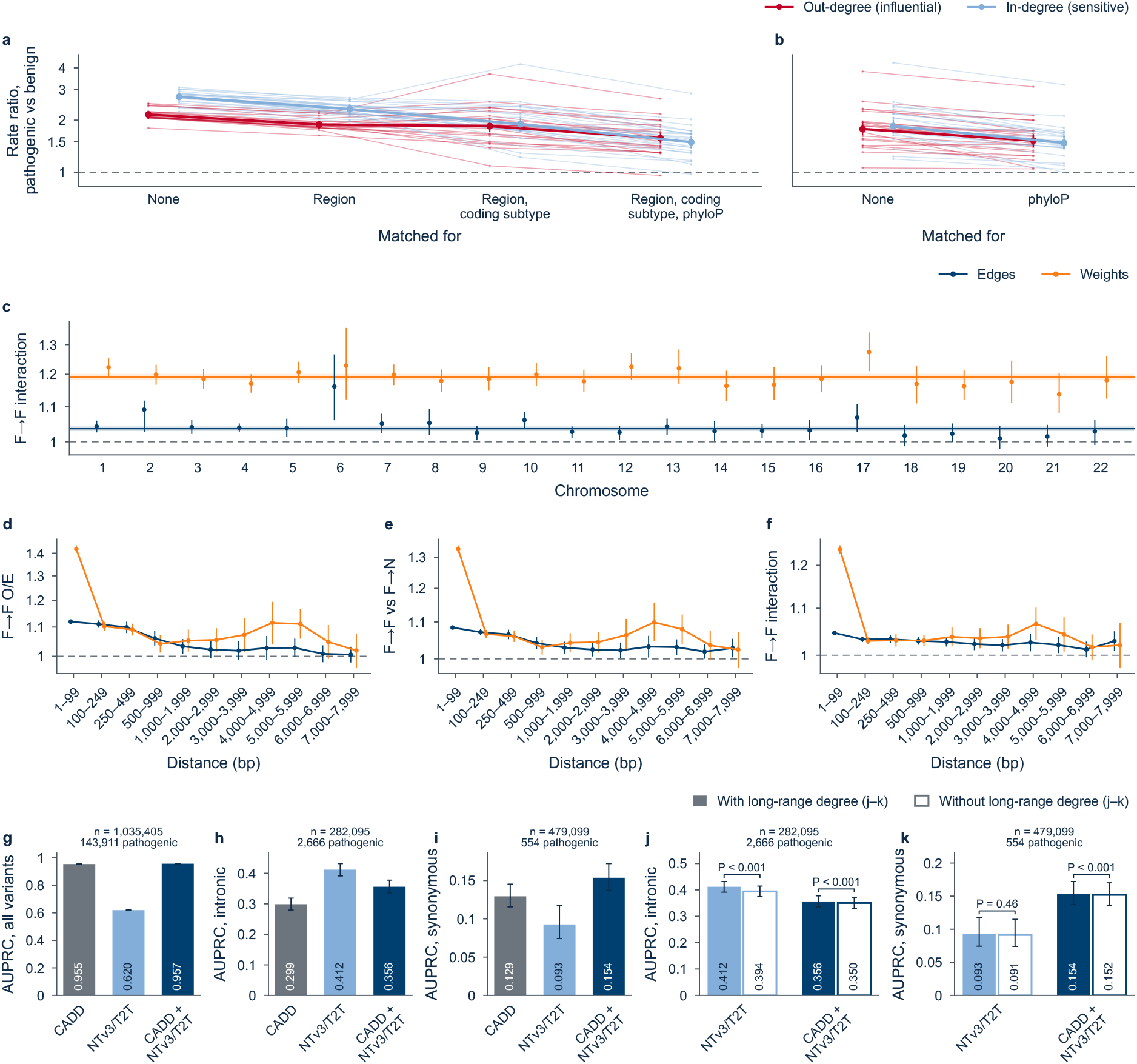
Nucleotide dependencies reflect human genetic variation and pathogenicity. **a,b**, Relative outgoing (red) and incoming (blue) dependency counts at pathogenic versus benign ClinVar positions, with successive adjustment for genomic region, coding consequence and evolutionary conservation (a), or restricted to missense positions with and without phyloP adjustment (b). Thin lines show individual autosomes; thick lines indicate pooled estimates with 95% confidence intervals (CIs). **c**, Enrichment of dependencies between frequent gnomAD variant positions (F; allele frequency >5%) beyond independent source and target effects, measured by edge counts (1.04-fold) and weights (1.19-fold). Points represent autosomes; horizontal lines indicate pooled estimates with 95% CIs. **d–f**, Distance-dependent enrichment of frequent-to-frequent dependencies: observed-to-expected ratios (d), enrichment relative to frequent-to-less-frequent dependencies (e), and interaction beyond independent source and target effects (f). Blue, edge counts; orange, edge weights. The terminal 8,000–8,191-bp bin is omitted. **g–i**, Pathogenicity prediction performance, measured by area under the precision–recall curve (AUPRC), for CADD v1.7, NTv3/T2T and their combination, across all ClinVar variants (g; n = 1,035,405), intronic variants (h; n = 282,095) and synonymous variants (i; n = 479,099). **j,k**, Ablation analyses for intronic (j) and synonymous (k) variants, comparing models with and without long-range out-degree (outgoing dependencies spanning ≥2 kb). Error bars indicate 95% CIs; P values indicate differences between models. Dashed reference lines in a–f indicate no enrichment.

We next examined common population variants in gnomAD^38,39^ (allele frequency >5%) across the autosomes (**Supplementary Fig. 8**). In contrast to pathogenic variants, common positions showed only marginally increased outgoing (1.015-fold) and incoming connectivity (1.018-fold). Dependencies between common positions were slightly more frequent than those connecting common to less frequent positions (1.053-fold), but showed little additional enrichment after accounting for the properties of their endpoints (1.036-fold). Their weights were more strongly enriched (1.249-fold relative to common-to-less-frequent dependencies; 1.19-fold beyond independent endpoint effects), particularly at distances below 2 kb (1.20-fold versus 1.05-fold at larger distances; **Fig. 7c–f**). Thus, common variation also exhibits relational organization, but with considerably weaker connectivity effects.

Finally, we asked whether nucleotide dependencies provide information beyond established computational pathogenicity scores. We combined long-range connectivity (outgoing dependencies spanning at least 2 kb) with NTv3 sequence-to-function and genomic feature predictions, and evaluated the resulting NTv3/T2T score alongside CADD v1.7^40^ in 1,035,405 ClinVar variants. Across all variants, CADD performed strongly (AUPRC 0.955), whereas NTv3/T2T reached 0.620. Combining both scores yielded only a small improvement (0.957; **Fig. 7g**). More pronounced differences emerged in variant classes that remain particularly challenging to interpret. For intronic variants, an important category of non-coding variation, NTv3/T2T substantially outperformed CADD (AUPRC 0.412 versus 0.299), while the combined model reached 0.356 (**Fig. 7h**). Additional value was also observed for synonymous variants, which do not alter the encoded amino acid. Although NTv3/T2T alone performed less well than CADD (0.093 versus 0.129), combining both scores improved AUPRC to 0.154, an increase of approximately 19% over CADD alone (**Fig. 7i**). Removing long-range connectivity reduced predictive performance, particularly for intronic variants, and also produced smaller but significant losses in the combined model (**Fig. 7j,k**; **Supplementary Fig. 9-10**). These findings indicate that nucleotide-dependency features contribute information beyond established variant scores, particularly for intronic and, in combination with existing scores, synonymous variants. Together, these findings link relational genome organization to human genetic variation and suggest that nucleotide dependencies may provide complementary information for variant interpretation, particularly where conventional approaches remain limited.

### Relational patterns in genetic disorders

To explore the potential relevance of relational genome architecture to human disease, we examined several loci associated with monogenic disorders (**Fig. 8**). The α-globin cluster, essential for haemoglobin production, contains *HBA1* and *HBA2*, which encode identical proteins, and the related, developmentally regulated *HBQ1* gene. Despite their proximity and sequence homology, these loci exhibit strong yet markedly asymmetric nucleotide dependencies, including directional relationships between *HBA1* and its neighbouring *HBQ1* gene (**Fig. 8a-c**). These observations reveal distinct relational architectures among closely related genes at a locus where genomic rearrangements are a well-established cause of α-thalassaemia. Whether these relational differences contribute to the distinct functional properties of these genes remains to be established.

**Fig. 8.**
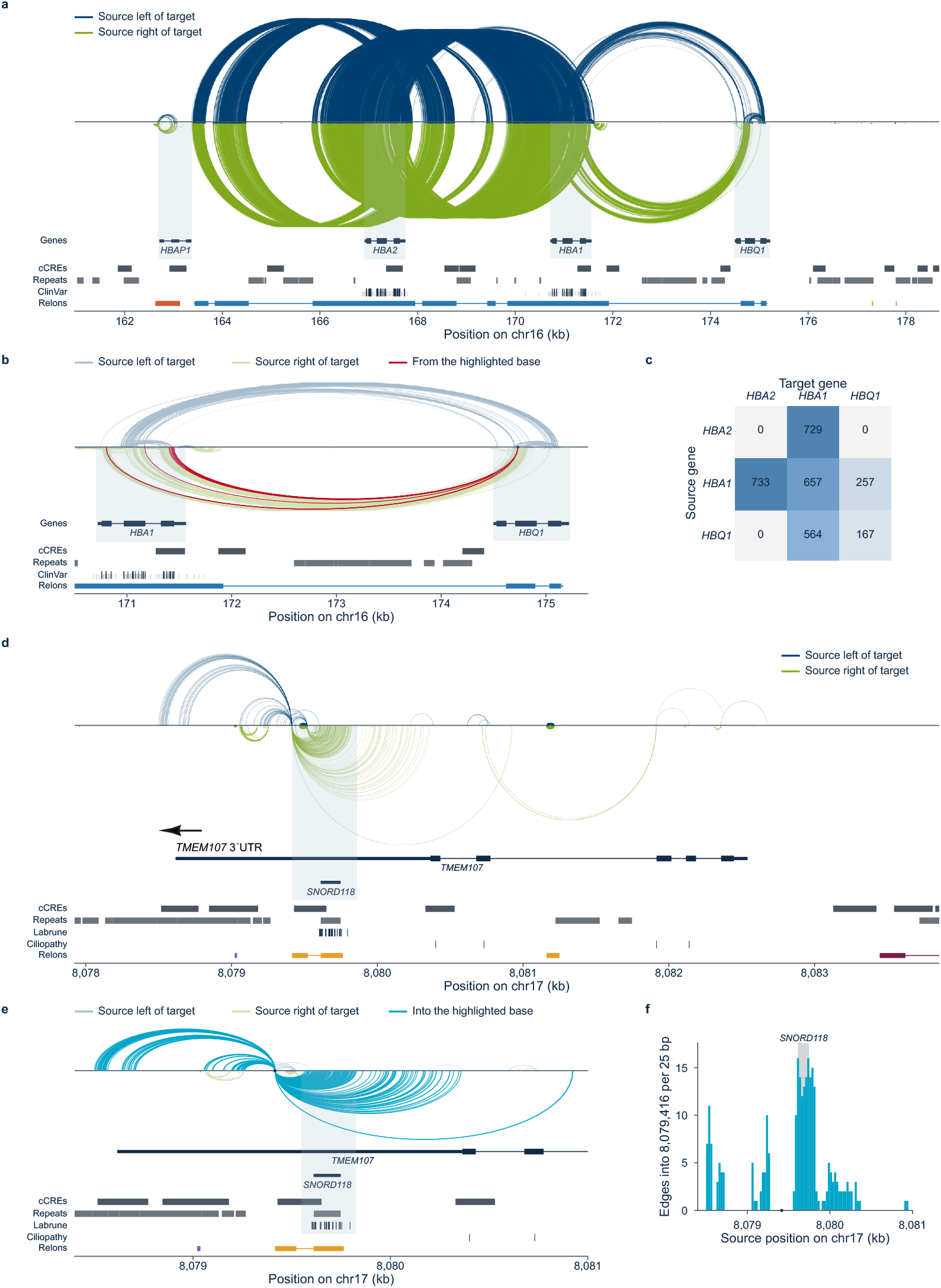
Directional nucleotide dependencies at monogenic disease loci. **a**, Strong dependencies (top 0.1% per genomic separation and cytoband class; 12,272 edges) across the α-globin cluster (T2T-CHM13v2.0, chr16:160,976– 178,674). Blue arcs indicate dependencies towards higher genomic coordinates, green arcs the reverse; opacity reflects edge strength. Edges connect perturbed source positions to responding targets. Tracks show genes, ENCODE cCREs, repeats, ClinVar variants (pathogenic variants highlighted) and Relons. **b**, *HBA1–HBQ1* region, highlighting 14 outgoing dependencies from *HBQ1* position chr16:174,734 to HBA1 (red), with no incoming strong edges. **c**, Directed dependency counts between globin genes reveal contrasting architectures: *HBA2* and *HBA1* show nearly symmetric connectivity (729 versus 733 edges), whereas *HBQ1–HBA1* relationships are markedly asymmetric (564 versus 257). Within-gene dependencies also differ (*HBA1*, 657; *HBA2*, 0), and no strong *HBA2–HBQ1* dependencies were detected. **d**, Dependency architecture at the overlapping *TMEM107–SNORD118* locus (chr17:8,077,924–8,083,851; 538 edges), with pathogenic ClinVar variants associated with Labrune syndrome and ciliopathies. **e**, Enlarged view of the *TMEM107* 3′ UTR, highlighting 263 incoming dependencies (cyan) at chr17:8,079,416, the beginning of a periodic Relon, with no outgoing strong edges. **f**, Distribution of source positions for these 263 dependencies (25-bp bins), showing their concentration around *SNORD118*.

A second example illustrates relational organization within overlapping disease-associated genes. Annotation-free inspection of the *TMEM107* locus that is associated with autosomal-recessive ciliopathies^41^ revealed a prominent periodic Relon within its 3′ UTR, including a position with 263 incoming but no outgoing dependencies (chr17:8,079,416). The strongest incoming dependencies originated from a region containing pathogenic ClinVar variants associated with autosomal-recessive leukoencephalopathy with brain calcifications and cysts (Labrune syndrome)^42^. Restoring genomic annotations revealed that this region corresponds to *SNORD118*, which encodes the U8 snoRNA and is embedded within the 3′ UTR of *TMEM107*. Thus, the relational pattern highlights a distinct sequence unit within overlapping genes associated with different monogenic disorders (**Fig. 8d,e**).

Finally, we examined an intronic *RFX3* variant (chr9:3,267,537; NM_001282116.2) identified in a patient with a neurodevelopmental disorder. Pathogenic *RFX3* variants have been associated with intellectual disability, developmental delay and autism spectrum disorder. The variant is located deep within the intron between exons 12 and 13 of *RFX3* and is embedded in an extensive network of directed nucleotide dependencies extending towards the flanking exons, suggesting a possible relationship to splicing (**Supplementary Fig. 11**). Subsequent analysis of patient-derived blood RNA indeed demonstrated aberrant splicing with inclusion of a 125-nucleotide pseudoexon, establishing the molecular consequence of the variant and supporting the diagnosis. Together, these examples illustrate how relational maps provide complementary views of disease-associated genomic regions and motivate further investigation of their potential utility in interpreting pathogenic variants in genetic disorders.

## Discussion

Complex systems are characterized not only by their individual components but also by the relationships between them. The human genome exemplifies this principle. Although its linear sequence, epigenetic modifications, regulatory interactions and three-dimensional organization are increasingly well characterized, a fundamental question remains: how do individual nucleotides depend on one another? In regulatory motifs or splice signals, for example, the significance of one nucleotide depends on its sequence context. Such relationships can be quantified computationally by asking whether substituting one nucleotide changes a genomic model’s prediction at another position. Whether these local rules are applicable to a higher-order architecture across the genome has remained unknown. Here, we combined genomic foundation models with large-scale GPU computing to systematically interrogate all 3.12 billion positions of the complete telomere-to-telomere human reference genome. We find that nucleotide dependencies follow characteristic rules of genomic distance, directionality and sequence context, repeatedly assembling into few distinct architectures, which we term Relons. These findings suggest a complementary perspective on genomic organization alongside established linear, functional and three-dimensional representations^8–15^. The resulting genome-wide relational landscape, the Relome, provides an accessible framework for exploring this organization at nucleotide resolution.

Several independent observations connect this relational landscape to established biological principles, demonstrating that model-derived nucleotide dependencies reflect biological organization rather than computational patterns alone. Dependencies preferentially connect evolutionarily conserved positions, even after accounting for genomic distance and the independent conservation of their endpoints^34^. This association becomes particularly pronounced at larger genomic separations, indicating that evolutionary constraint is reflected not only in individual nucleotides but also in their relationships. Relons further capture characteristic features of gene organization, including exon-to-exon connectivity and the three-base periodicity of coding sequence, without requiring gene annotations for their detection. Despite substantial regional differences in Relon distribution, the relative abundance of Relon classes remains remarkably consistent across chromosomes. Moreover, recurrent architectures extend into regions lacking current genomic annotation, defining thousands of Dark Relons. Together, these findings reveal a structured relational landscape that reflects established biological organization while extending beyond the boundaries of annotated genomic elements. This opens opportunities to investigate how genomic sequence is organized in regions where established annotations provide limited insight, raising the possibility that some of these regions harbour previously unrecognized biological functions.

The association of relational organization with human genetic variation also opens opportunities for clinical genomics. Positions harbouring pathogenic ClinVar variants exhibit substantially increased connectivity compared with benign positions, even after adjustment for coding consequence and evolutionary conservation. Associations with common and complex-trait variants are more modest, suggesting that relational connectivity captures different aspects of human genetic variation. Much of the preferential connectivity between pathogenic positions reflects their higher overall connectivity rather than a distinct disease-specific network. These findings show that positions with pathogenic variants exhibit increased relational connectivity compared to benign positions, even after adjustment for coding consequence and evolutionary conservation^38,43^. Additionally, we have demonstrated that network derived features can be integrated into a pathogenicity prediction model and encapsulates information that synergizes with sequence to function predictions. A relational perspective may be especially informative for non-coding and splice-altering variants, overlapping genes, and complex genomic regions where conventional annotation provides an incomplete picture. The α-globin (*HBA1, HBA2*), *SNORD118*, and *RFX3* examples illustrate these possibilities at clinically relevant loci, including a variant with an independently established splicing consequence. By placing individual variants within their broader nucleotide-dependency context, the Relome provides a foundation for exploring how relational features might contribute to future approaches to variant prioritization and molecular diagnosis.

Nevertheless, nucleotide dependencies reflect model responses to *in silico* perturbations rather than experimentally established molecular interactions, and their direction does not establish biological causation. Sequence homology, repetitive elements and model-specific properties may influence the observed patterns, while Relon detection depends on the applied thresholds and background models. However, all Relon classes were also recovered under an alternative filtering strategy excluding dependencies involving annotated repeats and non-B DNA motifs, demonstrating that recurrent architectures are not confined to these sequence features. Notably, the pronounced enrichment of non-B DNA motifs in specific Relon classes is intriguing, as these sequences can adopt alternative DNA conformations. Whether relational patterns also reflect aspects of DNA structural organization, rather than the underlying sequence properties alone, seems plausible but remains to be explored. Relons thus provide a systematic description of recurrent nucleotide-dependency architectures, opening the way to targeted functional studies. Comparisons across models, species and human haplotypes, together with experimental perturbations, will help establish which patterns correspond to biological interactions or constraints^44,45^. Emerging long-context genomic models may further extend this framework beyond the local sequence windows examined here^46^.

Beyond genome annotation and clinical interpretation, relational architecture also raises questions about the evolutionary forces shaping genome organization. Evolution changes nucleotide sequences, yet biological functions frequently depend on combinations of positions rather than individual bases. Compensatory substitutions, for example, can preserve RNA structures, while regulatory elements may retain their activity despite substantial sequence divergence^47^. This raises the possibility that evolution may preserve not only individual nucleotides or sequence motifs, but also particular configurations of dependencies between them. Conversely, the reorganization of these relationships during duplication, divergence and adaptation could create opportunities for functional diversification. Such processes might contribute to two defining properties of complex biological systems: robustness to perturbation and the capacity to evolve new functions^48,49^. The sequence-independent classification of Relon geometries provides a framework for investigating whether relational architectures are themselves conserved or reorganized across evolutionary timescales. This suggests the intriguing possibility that genome evolution involves not only changes in nucleotide sequence, but also the preservation and reorganization of the relationships encoded within it.

Taken together, our study provides evidence for a recurrent, genome-wide organization of nucleotide dependencies, establishing a framework for exploring the human genome as a complex relational system.

## Methods

### Inference utilizing NTV3

#### Creation of dataset

The T2T reference genome (CHM13v2.0)^8^ was obtained from NCBI. For the nucleotide dependency calculations, the genome was chunked into 761,065 intervals sized 8,192 bases with a 4,096 bases overlap to avoid boundary neglections, which was performed utilizing Pysam^50^. For the sequence to function dataset, again Pysam was utilized, to create genomic intervals of size 8,192 that were centered on the position of interest, at which the to-be-perturbed nucleotide was located. The created windows were subsequentially handled using Pandas^51^ and Numpy^52^.

#### Inference runs

For the inference of the nucleotide dependency maps the NTV3 pre-model (650M version) was utilized, receiving input length windows of 8,192 bases, the model being set to evaluation mode. The resulting logits per sequence were extracted and transformed into nucleotide dependency maps, as described previously^30^. The sequence-to-function and genomic tracks predictions were performed utilizing the NTv3 post-model (650M version), also receiving (variant wise) input length windows of 8,192 bases, while the model was also set to evaluation mode. Both models were downloaded from Huggingface and utilized using Huggingfaces Transformers library^53^ and Pytorch^54^. For the genomic element and sequence to function track predictions, at first a log2 fold change per track was calculated between the variant track and the reference track. Subsequentially the track fold change was summarized to a max score per track, as described by:

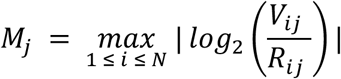

#### Prefiltering utilizing nucleotide dependency maps generated from randomized nucleotide sequences

Nucleotide dependencies were derived by analysing character probabilities from the NTV3 model (cite). This lead, to the situation that every node is related to every other node with at least a small weight. Due to performance reasons and for noise control we applied a prefilter step on every computed nucleotide dependency map. For this we additionally calculated 140 nucleotide dependency maps for random nucleotide sequences, which were created using the python random package. Subsequentially for the resulting 8,192 sized random windows, a binning approach was performed and edges were sorted into the bins depending on their distance. Across the resulting bins the 99th quantile was calculated and every edge was discarded that had lesser weight than this threshold.

### 1. Enrichment analysis across PhyloP

#### Data preparation

Base-wise PhyloP scores, obtained from UCSC^55^, (phyloP100way^34^ originally available for HG8) were lifted to the T2T assembly with Crossmap^56^. A position was classed as conserved (C) if the phyloP was greater than 2.7 and as not conserved (N) otherwise. Next to the threshold of 2.7, the analysis was repeated with thresholds of 1.5, 2.0 and 4.0. At first, every chromosome was analysed separately. Edges were read from the overlapping genomic windows. Edges with non-finite weights, self-loops and edges with an endpoint lacking a phyloP score were dropped. Edges reported in more than one overlapping window were merged into a single edge whose weight was the mean of the duplicates. Each edge was assigned one of four pair types according to the conservation of its source and target (C->C, C->N, N->C, N->N), and its genomic distance was assigned to one of 12 bins (1-99, 100-249, 250-499, 500-999 bp), then 1,000 bins up to 8,000-8,191 bp.

#### Calculating distance-matched expected values

To account for the potential clustering of conserved positions, expected values were derived from the candidate pairs available at each distance. A position was labelled a candidate position, if it either had at least one edge to a position with a PhyloP score or lay between two such positions no more than 1,000 bp apart. Subsequentially, the chromosome was divided into 500-kb blocks. Within each block b and for each signed offset d between source and target, we counted the candidate pairs P_b(d) and the candidate pairs of each pair type P_b,t(d), using fast Fourier transform-based^57^ cross-correlation of position indicator tracks. The expected number of edges of type t was:

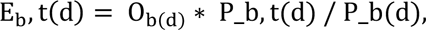

where O_b(d) is the observed number of edges at offset d in block b. Observed edges at each distance were therefore distributed over the pair types in proportion of the pairs of each type that were available at that distance. The expected values summed over the four types equal the observed edges. Expected edge weights were calculated in the same way, from the summed observed weight at each offset. Observed and expected values were summed over offsets within each distance bin and over blocks.

#### Calculating observed expected ratios

For each pair type, distance bin and statistic (edge count or edge weight), we computed the ratio of observed to expected values (O/E). From these we derived:

1. O/E C->C, the enrichment of connections between conserved positions relative to any pair at the same distance.
2. The ratio: (O/E C->C) / (O/E C->N), which measures, among conserved sources, the preference for conserved over non-conserved targets at the same distance.
3. The interaction: (O/E C->C * O/E N->N) / (O/E C->N * O/E N->), which measures enrichment of C->C pairs beyond the separate effects of a conserved source and a conserved target, on the risk-ratio scale.

Confidence intervals were obtained by a block bootstrap. The 500-kb blocks were resampled with replacement (10,000 replicates), and each ratio was recomputed from the resampled sums of observed and expected values. 95% confidence intervals were the 2.5th and 97.5th percentiles of the bootstrap distribution. Two-sided p-values were computed from the log estimate divided by the bootstrap standard deviation of the log ratio, using the normal distribution. Estimates were set to missing when fewer than 90% of bootstrap replicates were finite. P-values were adjusted across distance bins with the Benjamini–Hochberg^58^ procedure, separately for each threshold, statistic and metric.

#### Combining results across chromosomes

Finally, the individual results for the 22 autosomes were combined by a meta-analysis. Standard errors were recovered from the bootstrap confidence intervals as (log upper − log lower) / (2 × 1.96). The inverse-variance fixed-effect estimate was used as the main summary. DerSimonian–Laird random-effects estimates^59^, Cochran’s Q^60^ and I² ^61^ were computed to assess heterogeneity. For O/E metrics, the pooled ratio of summed observed to summed expected values across chromosomes is additionally reported. P-values were adjusted across distance bins with the Benjamini–Hochberg procedure.

### 2. Enrichment analysis across ClinVar

#### Data preparation and descriptive analysis

The lifted ClinVar release 20220313 was obtained from the official T2T-CHM13v.20 Github. The resulting release was filtered for likely pathogenic (P), pathogenic (P), likely benign (B) and benign (B) entries. Additionally, a star-based filter was performed, keeping only variants with at least one-star, while also dropping conflicting entries. For the analysis only positions at which only pathogenic or only benign variants occurred. The dependency network was restricted to edges in which both source and target were ClinVar positions classified as pathogenic or benign, so that every edge was of type P→P, P→B, B→P or B→B. Edges touching an unlabelled or excluded positions were excluded. Repeated edges between the same ordered pair of positions (due to the overlapping windows) were merged into a single edge whose weight was the mean of the duplicates. Subsequentially, we computed the fraction of outgoing edges that reached a pathogenic target, both by edge count and by summed edge weight. Enrichment was summarised as the risk ratio of these fractions (pathogenic over benign positions).

#### Node-label permutation tests

Next, we performed a permutation test. The test statistic was the fraction of edges from pathogenic sources that ended in a pathogenic target, computed by edge count and edge weight. Null distributions were generated by permuting node labels among the positions present in the analysed network, keeping network topology and edge weights fixed. Two null models were utilized, (i) an unrestricted label permutation, and (ii) a degree-preserving permutation in which labels were shuffled only within strata of nodes with similar out- and in-degree. For the stratified null, out- and in-degree were each divided into q rank-based quantile bins, with q = min(5, ⌊√(n/10)]) for n nodes, giving up to 25 strata. Each test used 5,000 permutations. One-sided p-values were computed as (1 + number of permutations with a statistic ≥ the observed value) / (1 + number of permutations). Evaluation metric were the fold enrichment (observed / mean of the null), z-score, and the 2.5^th^-97.5^th^ percentiles of the null distribution. Additionally, the permutation tests were repeated on subnetworks obtained by (i) removing the top 1% and top 5% of edges by weight, (ii) splitting edges into tertiles of the edge distance provided with the network, and (iii) separating reciprocal from non-reciprocal edges. Labels were permuted within each subnetwork. P-values were corrected for multiple testing with the Benjamini–Hochberg procedure, separately for each null model and each statistic (count and weight).

#### Distance-matched candidate pair models

To account for the genomic clustering of pathogenic variants, we enumerated all ordered pairs of network positions whose genomic distance did not exceed the largest observed edge distance (candidate pairs) and coded each pair as connected (an observed edge) or unconnected. Genomic distance was separated into 20 quantile bins of the observed edge distances, and into tertiles (short, medium, long).

As descriptive checks we report, by distance tertile, the fraction of candidate partners that were pathogenic for pathogenic and for benign sources, and the number of connections per 1,000 candidate pairs for each pair type (P→P, P→B, B→P, B→B). Connection was modelled with generalised linear models^62^:

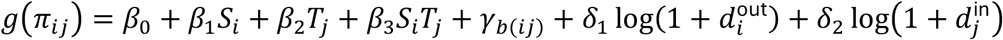

*π_i_*_j_ is the probability that source position i is connected to target position j, and g is the link function. *S_i_*and *T*_j_indicate whether the source and the target are pathogenic (1) or benign (0), *γ_b_*_(*i*j)_is a fixed effect for the distance for the distance bin b of the pair, and 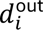 and 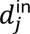 are the observed out-degree of the source and in-degree of the target. The interaction *β*_3_ tests whether pathogenic-to-pathogenic pairs connect more often than expected from the separate effects of a pathogenic source (*β*_1_) and a pathogenic target (*β*_2_). Models were fitted with a logit link g(π) = log[π / (1 − π)] giving odds ratios, and with a log link using Poisson regression, g(π) = log π, giving risk ratios. Each link was fitted with and without the degree terms (*δ*_1_, *δ*_2_). The logit model with degree adjustment was the main specification. Standard errors were clustered by source position. For each specification, the model without the interaction term was used to compute the ratio of observed to predicted P→P edges. Distance-specific interactions were estimated by replacing *β*_3_ with separate P→P terms for the short, medium and long distance tertiles, and heterogeneity across tertiles was assessed with a Wald test^63^.

Among connected pairs, edge weight was modelled by ordinary least squares on log(w + ε), with ε equal to half the smallest positive weight, adjusted for distance bin with and without the degree terms. We fitted (i) P→P edges against all other edges and (ii) the P→P interaction beyond pathogenic-source and pathogenic-target main effects. Exponentiated coefficients were reported as geometric mean weight ratios, with standard errors clustered by source position.

#### Evaluating whether Pathogenic Positions are dependency hubs

For every candidate pair, a label-free expected connection probability was set to the observed connection rate in its distance bin, and an expected weight to that probability multiplied by the mean weight of observed edges in the bin. For each position we summed observed and expected values over its candidate pairs to obtain out-degree and out-strength (as source) and in-degree and in-strength (as target). The same quantities were also computed using benign partners only.

Each metric was compared between pathogenic and benign positions with a Poisson generalised linear model^64^, observed ∼ pathogenic, with log(expected) as an offset, so that the exponentiated coefficient is the ratio of observed-to-expected rates for pathogenic versus benign positions. For non-integer strengths the model was used as a quasi-likelihood estimator. Standard errors were clustered by 10 kb genomic blocks. Median observed/expected ratios are reported per class.

At the pair level, connection was modelled as connected ∼ P_source + P_target + distance bin, with logit and log links, and edge weight among connected pairs as log(w + ε) ∼ P_source + P_target + distance bin (ordinary least squares). Standard errors were clustered by the 10 kb block of the source position.

#### Adjustment for genomic region and coding subtype

Each position was assigned a genomic region from the ClinVar molecular consequence annotation (obtained from the 20220313 release VCF), these being: splice related (“splice_acceptor_variant” and “splice_donor_variant”), coding related (“missense”, “synonymous”, “nonsense” and “stop_lost”), UTR related (“5 prime UTR”, “3 prime UTR”), “non-coding transcript”, “intronic”, “upstream” or “downstream” variant. Where variants at one position had multiple consequences, the priority of assignment followed the stated order. Positions without annotation or with unmapped consequences were assigned to an “unknown” category.

The position-level hub models were refitted with region as a categorical covariate (reference: intron). The share of the effect remaining after adjustment was computed as

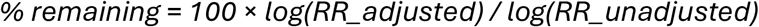

Region effects on out- and in-degree were estimated from the same adjusted models. Pathogenic and benign positions were also compared within each region separately, excluding the unknown category, for regions with at least 10 positions of each class.

Coding positions were further split by the variants observed at the position, in order of precedence: nonsense or stop-lost, missense, synonymous only, and other. The hub models were refitted with this finer classification as covariate and, where at least 10 positions of each class were available, within each coding subtype.

#### Adjustment for evolutionary conservation

Each position was annotated with its PhyloP score. Positions without a mappable PhyloP score were dropped. Subsequentially, the position-level hub models were fitted (i) without adjustment, (ii) with phyloP modelled as a natural cubic regression spline with 4 degrees of freedom, (iii) with the spline plus the finer region classification, and, among missense positions with at least 10 positions of each class, (iv) without and (v) with the phyloP spline. All models in this analysis used only positions with a phyloP score. The percentage of the log rate ratio remaining was computed as above, relative to the matching unadjusted model. The association of conservation itself with out- and in-degree was estimated per standard deviation of phyloP, entered linearly and adjusted for pathogenicity. Standard errors were clustered by 10 kb genomic blocks.

#### Meta analysis summarizing the results

As previously introduced, each of the 22 autosomes was analysed independently, and chromosome-specific estimates were combined by a subsequent meta-analysis. Every ratio estimate reported with a 95% confidence interval was pooled. Standard errors were recovered from the confidence intervals as (log upper − log lower) / (2 × 1.96). We computed the inverse-variance fixed-effect estimate and the DerSimonian–Laird^59^ random-effects estimate, with 95% confidence intervals and two-sided p-values from the normal distribution; random-effects estimates are reported as the primary summary. Heterogeneity was quantified with Cochran’s Q^60^ (with its χ² p-value on k − 1 degrees of freedom), I² and τ². Strata that could be estimated on only some chromosomes, such as within-region comparisons requiring at least 10 pathogenic and 10 benign positions, were pooled over the chromosomes available. The permutation test results were exempt from this meta analysis.

### 3. Enrichment analysis across gnomAD

#### Dataset preparation and preprocessing

The provided gnomAD 3.1 VCF files, which were originally lifted over from hg38 by the T2T consortium were retrieved. Subsequentially, only variants were kept that had the “PASS” keyword in the filter column. At multi-allelic positions, the allele with the highest allele frequency (AF) was retained, giving one max AF per position. Furthermore, sites with a low call rate (defined as allele number less than 50% of the chromosome’s maximum) were dropped. Only gnomAD variant sites took part in the analysis, positions with no frequency were dopped. Positions with AF > 0.05 were classed as frequent (F) and all other variant site (AF ≤0.05) as less frequent (N). The analysis were repeated with AF thresholds of 0.01 and 0.1.

#### Distance-matched enrichment

Edge processing, candidate pairs, distance-matched expected values, the pair-type statistics (O/E F→F, F→F vs F→N and the interaction) and the block bootstrap were as described for the conservation analysis, with candidate positions restricted to variant sites. Each chromosome was analysed separately. Subsequentially, estimates from the 22 autosomes were combined on the log scale, with standard errors recovered from the bootstrap confidence intervals as (log upper − log lower) / (2 × 1.96). The DerSimonian–Laird^59^ random-effects estimate was used as the main summary, with heterogeneity quantified by Cochran’s Q^60^, I² and τ².

#### Sequence-to-function based pathogenicity prediction

Each variant from the autosomes of the 20260104 Clinvar release were annotated with 78 features: the long-range-degree (defined as spanning at least 2,000 basepairs), 59 sequence-to-function tracks and 18 gene feature tracks, both predicted from the NTv3 post model. To capture sequence-to-function predictions across distinct cellular contexts while minimizing confounding by assay availability, we selected H1, K562 and HepG2 cells, representing pluripotent, haematopoietic and hepatic states, respectively. We restricted the analysis to functional readouts available in all three cell types, comprising DNase accessibility, H3K4me1, H3K4me3, H3K36me3, H3K9me3, H3K27me3, CTCF and POLR2A. After the exclusion of missing features, 1,035,405 variants remained (143,911 pathogenic, 891,494 benign).

The NTv3/T2T score was obtained by L2-regularized logistic regression on standardized features (scikit-learn [version], default settings, maximum 2,000 iterations). A second logistic regression combined the standardized raw CADD score with the predicted probability of the NTv3/T2T model (CADD + NTv3/T2T). Both models were fitted only on the CADD training data; no ClinVar variant was used for model fitting. Models were trained on the CADD training set (CADD 1.7), comprising 13,625,521 simulated and 13,774,107 human-derived variants. Simulated variants served as the target class, following CADD’s training objective. Variants with missing values in the raw CADD score or any feature were excluded.

Molecular consequences were taken from the ClinVar MC annotation, and variants were grouped as:

- splice site: splice donor or acceptor variant
- loss of function: nonsense, stop-lost or initiator codon variant
- missense
- synonymous
- 5′ UTR
- 3′ UTR
- intronic

In a non-exclusive analysis, a variant was assigned to every class matching any of its annotated consequences. In an exclusive analysis, each variant was assigned to one class by the priority splice site > loss of function > missense > synonymous > 5′ UTR > 3′ UTR > intronic. Variants without a matching consequence formed an additional class. Classes with fewer than 50 variants were not evaluated.

Performance was measured as ROC-AUC and AUPRC, computed as average precision. 95% confidence intervals were obtained from 1,000 bootstrap resamples of variants, drawn with replacement separately within each class (percentile method). All models were evaluated on the same resamples, so differences between models are paired. Two-sided P values were computed from the bootstrap distribution of each difference (resolution 0.001).

To assess the contribution of long-range degree, both models were retrained without this feature. Training and evaluation variants were kept identical to the full model by applying the same exclusion criteria based on all 78 features.

### 4. Relon analysis Preprocessing

#### Stiching overlapping windows

Dependency maps were computed in windows of 8,192 bp that start every 4,096 bp, so each position lies in two windows. An edge *i* → j runs from a source position i, the position that is perturbed, to a target position j, the position whose prediction changes. Its separation is d = |j − i|. A pair closer than 4,096 bp can lie in both windows that overlap it. A pair 4,096 bp or more apart fits into one window only. We therefore merged the windows into one record per ordered pair. For each pair, we recorded the number n_ij_ of measured windows that contain both positions, and the number r_ij_ ≤ n_ij_ of these windows that reported the edge. From these, we computed the presence p_ij_ and the stitched log weight *l*_ij_:

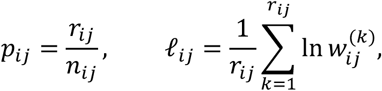

where 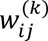 is the weight reported by the *k*-th window that reported the edge. *l_i_*_j_ is thus the natural logarithm of the geometric mean of the reported weights. The archive stores only edges that passed the low-confidence filter described earlier (Methods, *Prefiltering utilizing nucleotide dependency maps generated from randomized nucleotide sequences)*. A window that contains a pair but did not report its edge, because the edge was removed by this filter, counts as a measurement of zero in the presence. Otherwise, pairs closer than 4,096 bp would have two chances to pass the filter, and longer pairs only one. Pairs that no window reported (*r_i_*_j_ = 0) were not stored, but still count as pairs that could have been measured. Windows absent from the archive or unreadable were treated as not measured: they contribute neither edges nor pairs that could have been measured. Each pair was assigned to the 4,096 bp block that contains its lower position, min(*i*, *j*), and took the cytoband class^8^ at the midpoint of that block: gene-rich (gneg), gene-poor (gpos25 to gpos100), heterochromatin (gvar), rDNA (stalk) or centromere (acen).

#### Edge filters

Edge weights and the number of stored edges change strongly with separation and cytoband class. A single weight cut-off would therefore select mostly adjacent positions and repeat-rich regions. We instead selected strong edges relative to their own separation and cytoband class, with two filters.

*Strongest 0.1 %.* We selected strong edges within groups. A group *G* holds all stored pairs of one chromosome that share the same cytoband class *c* and the same exact separation *d*, with *d* ≥ 2. Within each group, we kept the pairs with the highest stitched log weight until they made up 0.1 % of the group. In this count, each pair contributes its presence *p_i_*_j_, not 1. A pair reported by every window that contains it counts as 1; a pair reported by only one of its two windows counts as 0.5. A pair therefore never counts twice. Presence is used only in this count: whether a pair is strong depends on its log weight alone. The threshold of a group is

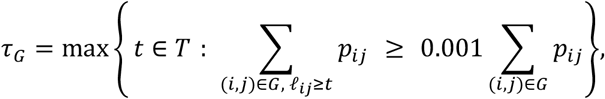

where *T* = {−3.60, −3.55, …, 3.40} is a grid of natural log weights in steps of 0.05. A pair is strong if *l_i_*_j_ ≥ *τ_G_*. Because the threshold lies on this grid, slightly more than 0.1 % of each group is kept (about 0.108 %). The 0.1 % is a share of the edges that passed the low-confidence filter, not of all possible pairs. Unless stated otherwise, strong edges are those of this filter.

*Strongest 1 % without repeats.* Repeats contribute many of the strongest edges. To see which structure remains without them, we built a second filter on repeat-free sequence. We masked every base annotated by RepeatMasker (v4.1.2p1.2022Apr14, T2T consortium release) and every base in one of the seven non-B DNA motif types of the T2T consortium annotation (G-quadruplex, Z-DNA, inverted, mirror, A-phased, short tandem and direct repeats). Bases of RepeatMasker’s RNA classes (rRNA, snRNA, srpRNA, scRNA, tRNA) and exons of small RNA genes in GENCODE^65^ (for example snRNA, snoRNA, miRNA; not lncRNA) were left unmasked, because they are RNA genes rather than copy-number artefacts. An edge was removed if its source or its target lies on a masked base. Masking came first: the threshold was then set on the remaining pairs, with the same rule as above and 1 % instead of 0.1 % (0.01 instead of 0.001 in *τ_G_*). Masked pairs were also removed from the pairs that could have been measured: they can hold no edge, so counting them would dilute the expected density of edges between the repeats. The two filters differ in two ways at once, the mask and the cut (1 % against 0.1 %).

### Strong-edge enrichment between element classes

This analysis uses the strongest 0.1 % filter. Each position was given one class per annotation: its cCRE class and its gene element. Positions outside any element formed a class of their own (no cCRE, intergenic). We compared cCRE classes as sources with gene elements as targets, and cCRE classes with cCRE classes. We grouped separations into five bands *b*: 2–15, 16–127, 128–1,023, 1,024–4,095 and 4,096–8,191 bp. For a source class *A*, a target class *B*, a cytoband class *c* and a band *b*, let *P*(*A*, *B*, *c*, *b*) be the stored pairs *i* → *j* with *i* in class *A*, *j* in class *B*, cytoband class *c* and *d* in band *b*. We counted all edges *N* and strong edges *S* in this set:

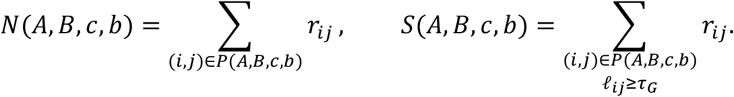

Each pair counts once for every window that reported it, so these counts are the same as those obtained by reading the windows one at a time. Strong edges and all edges are counted in the same way. The totals over all class pairs are *N*(*c*, *b*) = ∑*_A_*_,*B*_ *N* (*A*, *B*, *c*, *b*) and *S*(*c*, *b*) = ∑*_A_*_,*B*_ *S* (*A*, *B*, *c*, *b*). The enrichment compares the share of strong edges of one class pair with the share of strong edges among all edges in the same band and cytoband class:

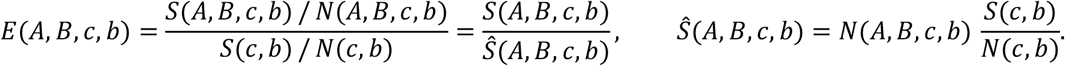

*S*^r^ is the number of strong edges expected if the class pair held the same share of strong edges as all edges in its band and cytoband class. *E* = 1 means no preference; *E* = 2 means twice and *E* = 0.5 half the expected number of strong edges. Strong edges were selected per chromosome, as described above, and all counts were then summed over the 24 chromosomes before any ratio was taken. Cells with *S*^r^ < 30 were not interpreted.

### 5. Relon detection heuristic and analysis

#### Expected strong edges

Relons were detected separately for each chromosome and each edge filter. For cytoband class *c* and separation *d*, the probability that a pair carries a strong edge is

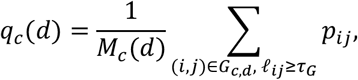

where *G_c_*_,*d*_ is the group of stored pairs in class *c* at separation *d* and *M_c_*(*d*) is the number of ordered pairs at separation *d*, with the lower position in class *c*, that a measured window contains. Under the repeat-free filter, masked pairs are not counted in *M_c_*(*d*). Detection worked on tiles, the 8,192 bp stretches covered by the windows. For each tile we built two matrices over its positions, with sources as rows and targets as columns. The observed matrix holds *O_i_*_j_ = *p_i_*_j_ for a strong edge and 0 otherwise. The expected matrix holds *X_i_*_j_ = *q_c_*(*d*), where *c* is the class of the lower position; it is 0 where no measured window contains the pair or where the pair is masked. Separations 0 and 1 hold no strong edges and are 0 in both matrices.

#### Box ratio, seeds and growth

We scanned both matrices with square boxes of *b* = 4, 16, 64, 128 and 256 bp, so that patterns of a few bases and of several kilobases are both captured. Boxes are centred on grid points *b*/4 apart, so neighbouring boxes overlap, and are cut off at the edge of the tile. For each box we summed the observed and expected values, *O_b_* and *X_b_*, and computed the log2 box ratio

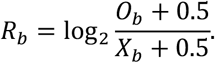

The pseudocount 0.5 keeps a box with a single stray edge, where almost none is expected, from scoring a thousandfold enrichment. A box is a seed if its summed presence is at least 3 (*O_b_* ≥ 3) and is at least eightfold enriched at *b* = 4 bp (*R_b_* ≥ 3) or at least four-fold at the other box sizes (*R_b_* ≥ 2). From each seed, a candidate grows over neighbouring grid points, touching at a side or a corner, as long as they are at least twofold enriched (*R_b_* ≥ 1) and hold an edge (*O_b_* > 0). A candidate is thus a connected set of such grid points that contains at least one seed. It is kept only if its strongest seed also reaches the chance maximum *θ_b_*_,*c*_ for its box size and for the cytoband class of its first base.

#### Null map and chance maximum

The chance maximum was calibrated on a simulated null map. The null map replaces the observed matrix by edges placed at random, while the expected matrix stays the same. Every cell is drawn independently: it has presence 1 with probability 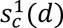, presence 0.5 with probability 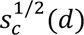, and no edge otherwise. Here 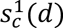 and 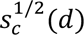 are the shares of measurable pairs in class *c* at separation *d* that carry a strong edge at presence 1 and 0.5. The mean of each cell is therefore its expected value, 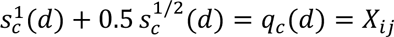 On average, the null map thus holds as many strong edges at each separation and in each cytoband class as the real map, but scattered at random, without blocks, stripes or hubs other than those that arise by chance. The edges are drawn fresh: no real edge is moved or reused, so a large Relon cannot raise its own threshold. The box ratio *R_b_* of the null map is computed as above, with the null map in place of *O* and the same *X*. To set the chance maximum, we collected the box ratios of all null boxes, separately for each box size *b* and cytoband class *c*, and took the far end of their distribution: *θ_b_*_,*c*_ is the lowest ratio that at most one in 10^7^ null boxes reaches or exceeds (ratios counted in bins 0.05 wide),

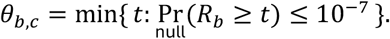

Where too few null boxes were drawn to resolve one in 10^7^, the highest null box was used instead.

#### Projection onto the genome

The area *A* of a candidate is the union of the boxes of its grid points, which reaches up to half a box beyond the pattern. We therefore trimmed its rows and columns: a row *i* was kept if it holds at least half an edge more than expected within the area, ∑_j: (*i*,j)∈*A*_ ( *O_i_*_j_ − *X_i_*_j_) ≥ 0.5, and likewise for columns. The edges of the candidate are the strong edges in the area that lie in kept rows and kept columns. Their source and target positions, split wherever two positions lie more than *b* apart, give the stretches of the candidate. Because tiles overlap by half, most candidates are found by two tiles. Each was kept by the tile whose central 4,096 bp contain the middle of its span.

#### Relon classes

Each candidate was given a class from the positions of its edges. Let *w_S_* and *w_T_* be the widths of the ranges of its source and target positions (largest minus smallest position, plus 1), *r* the Pearson correlation between source and target positions over its edges, and *f*_3_ the share of its edges at its three most frequent separations. The source and target ranges are *apart* if they do not overlap. The rules were applied in this order, and the first that holds gives the class: (1) source hub, 4*w_S_* ≤ *w_T_*; (2) target hub, 4*w_T_* ≤ *w_S_*; (3) anti-diagonal, *r* ≤ −0.5; (4) offset diagonal, apart and *r* ≥ 0.5; (5) island, apart; (6) periodic, *f*_3_ ≥ 0.5; (7) block, otherwise. The anti-diagonal is tested before the ranges must be apart, so an anti-diagonal can cross the diagonal.

#### Merge candidates into Relons

One structure is usually found as several candidates: at several box sizes, once for each direction of its edges, and in fragments. We merged these candidates into Relons in three steps.

First, we removed candidates that only thicken the diagonal, that is, that consist of edges between near neighbours rather than of a pattern. A candidate was removed if more than 60 % of its edges lie at separation 2 or 3, or if its reach is below 0.15. The reach is the 90th percentile of the candidate’s edge separations divided by the length of its span.

Second, we replaced coarse candidates by tighter ones. A large box reaches up to half a box beyond a pattern and picks up stray edges there. If a smaller candidate lies inside a larger one and holds at least 80 % of its edges, the larger candidate was replaced by the smaller one (by the smallest, if there are several). Inside means that at least 80 % of the smaller candidate’s bases lie in the larger one’s stretches and that it overlaps each of these stretches.

Third, we joined the candidates that belong together. Two candidates belong to the same Relon if their stretches overlap, or if at least one strong edge of the viewer tier runs between their stretches. This holds transitively: if A belongs with B and B with C, all three form one Relon. A Relon covers all stretches of its candidates. It takes the class and box size of its candidate with the most edges, or of the smaller box size on a tie.

#### Relon classes and genomic elements

For each Relon class *k* and genomic element *e* we compared the share of the class’s bases inside the element with the element’s share of all bases:

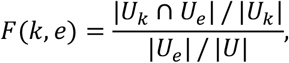

where *U* is the set of euchromatic bases (Giemsa bands gneg and gpos) of all 24 chromosomes, under the repeat-free filter only the unmasked ones, *U_k_* ⊆ *U* the bases in stretches of Relons of class *k*, and *U_e_* ⊆ *U* the bases inside element *e*. *U* includes the bases outside every Relon, so |*U_e_*|/|*U*| is the share a random euchromatic base would show. *F* = 1 means that the class lies on the element as often as a random euchromatic base. We restricted the count to euchromatin because, counted in bases, the few mega-base-long Relons of centromeres and heterochromatin would otherwise decide every class they fall into. Each base takes the class of the Relon it lies in. Counts were summed over the 24 chromosomes before the ratio was taken, and elements covering less than 0.5 % of *U* were not tested. The elements were RepeatMasker repeats (every class except rRNA); non-B DNA motifs (any of the seven types); gene parts from GENCODE v50, lifted from GRCh38 to T2T-CHM13 with liftOver, one per base in the order CDS, UTR, non-coding exon (any exon of a transcript without CDS), intron, and intergenic otherwise; ENCODE cCREs (SCREEN, lifted to T2T-CHM13)^10,66^, any class; and CpG islands (UCSC).

#### Exon Bridging

For each gene we chose one transcript with at least two exons: the MANE Select transcript for protein-coding genes and, for lncRNA genes, the transcript with the most exons (then the longest). Transcripts were taken in order of their start, skipping any that overlaps an already chosen transcript of the same gene type. The intron of a transcript is every base of its span outside its exons, so exons of other transcripts can count as intron, which makes the test conservative. An ordered pair of positions within one transcript belongs to one of four kinds *K*, by where its source and target lie: exon to another exon, exon to intron, intron to exon, and intron to intron; pairs within one exon were left out. For each kind and separation bin *b* (100–299, 300–999, 1,000–2,999 and 3,000–8,191 bp), the edge rate is

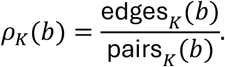

Here edges counts the strong edges whose two ends lie in stretches of the same Relon, and pairs counts all ordered pairs of that kind, the places where such an edge could lie. Rates are per pair rather than per edge because introns are much longer than exons. Exon bridging is

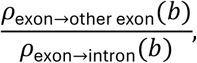

and the control is the same contrast from intron sources, *ρ*_intron→exon_(*b*)/*ρ*_intron→intron_(*b*), which would be as high if exons attracted edges from anywhere. Under the repeat-free filter, masked bases were removed from the pairs as from the edges. Edges and pairs were summed over the 24 chromosomes before the rates were taken.

#### Codon period in periodic candidates

We analysed the periods on candidates rather than on Relons, because only a candidate’s own edges carry its period, and a Relon takes its class from its largest candidate. We used the periodic candidates the Relons are merged from. A candidate counts as coding if at least half of its stretch bases are CDS. For each group *g*, coding or other, we collected the strong edges of the viewer tier with both ends in the stretches of a candidate of the group, each edge once per group even if several candidates hold it, and counted them per separation, *n*_g_(*d*), for separations from 4 to 63 bp. The share of separation *d* is

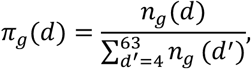

and the share at a multiple of 3 is the share of edges at 6, 9, 12, …, 63 bp:

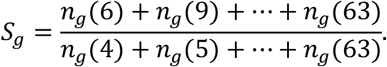

Of the 60 separations from 4 to 63 bp, 20 are multiples of 3, so evenly spread edges would give *S*_g_ = 1/3. Edge density falls with separation, however, so edges are not evenly spread even without a period. The baseline is therefore the other group, not a third.

Separations 2 and 3 were left out because they reflect closeness, not a period. Every candidate on the diagonal has many edges between near neighbours, about a third of the edges of every periodic candidate lie at 2 or 3 bp, whether or not the candidate has a period of 3. With separation 2 included, these edges would dominate the denominator of *S*_g_ and pull it down; with separation 3 included, they would count as a multiple of 3 and push it up. In either case, *S*_g_ would mostly measure how many edges lie between near neighbours, not the period. Leaving out 3 bp loses the closest codon distance, but the period still shows at 6, 9, 12 bp and beyond.

#### Compute systems and software

The supercomputers at the Jülich Supercomputing Centre (JSC) were used for this study to perform the calculations within a short time frame. All GPU-accelerated computations were performed on JUPITER, the first European exascale supercomputer and the world’s fastest NVIDIA-accelerated supercomputer publicly available for scientific research. JUPITER has 5,884 nodes, each with four NVIDIA GH200 superchips connected via a highspeed InfiniBand network in a Dragonfly+ topology^67^. The CPU-only computations were performed on the smaller supercomputers JUWELS^68^ and JURECA^69^, which have CPU-only nodes equipped with Intel Xeon Platinum 8168 and AMD EPYC 7742 processors, respectively. Analyses were performed in Python. Statistical analysis were performed using NumPy^52^, Pandas^70^, SciPy^71^ and Statsmodels^72^. The random seed was fixed (42) for all permutations.

#### Use of generative AI

OpenAI ChatGPT and Anthropic Claude were used to assist with language editing and text organization, as well as code development, debugging and computational data analysis. All AI-assisted code, analyses and scientific interpretations were critically reviewed and validated by the authors, who take full responsibility for the results and conclusions.

## Code availability

A Repository of the code is attached as supplementary file to the manuscript and will be published upon publication.

## Data availability

All preexisting datasets, utilized in this study are openly available at their corresponding repositories, these being: 1. Phylop100way from UCSC (https://hgdown-load.cse.ucsc.edu/goldenpath/hg38/phyloP100way/) 2. ClinVar 20220313 releases lifted to T2T by the T2T consortium (https://github.com/marbl/CHM13/blob/master/README.md) 3. gnomAD 3.1 releases lifted to T2T by the T2T consortium (https://github.com/marbl/CHM13/blob/master/README.md) 4. T2T assembly from NCBI (https://www.ncbi.nlm.nih.gov/datasets/genome/GCF_009914755.1/) 5. Resources utilized to construct genomic element masks for the Relome Browser from NCBI (https://ftp.ncbi.nlm.nih.gov/genomes/all/GCF/009/914/755/GCF_009914755.1_T2T-CHM13v2.0/) 6. Cyto Bands for T2T obtained from the T2T consortium (https://hgdownload.soe.ucsc.edu/gbdb/hs1/cytoBandMapped/cytoBandMapped.bb) 7. Gencode v50 Release (https://ftp.ebi.ac.uk/pub/databases/gencode/Gencode_human/release_50/gencode.v50.annotation.gtf.gz) 8. Gencode v50 tRNAS (https://ftp.ebi.ac.uk/pub/data-bases/gencode/Gencode_human/release_50/gencode.v50.tRNAs.gtf.gz) 9. Gencode Phase 2 Ribo-seq ORFs (https://ftp.ebi.ac.uk/pub/databases/gencode/riboseq_orfs/phase2/Ribo-seq_ORFs.comprehensive.bed) 10. Encode screen elements (https://screen.wenglab.org/downloads) 11. Repeat Masker v4.1.2 lifted from the T2T consortium (https://s3-us-west-2.amazonaws.com/human-pangenomics/T2T/CHM13/assemblies/annotation/chm13v2.0_Repeat-Masker_4.1.2p1.2022Apr14.bed) 12. Non-B DNA annotations from the T2T consortium (https://s3-us-west-2.amazonaws.com/human-pangenomics/T2T/CHM13/assemblies/annotation/nonB/) 13. UCSC simpleRepeat files (https://hgdown-load.soe.ucsc.edu/gbdb/hs1/bbi/simpleRepeat.bb) 14. UCSC cpgIsland files (https://hgdownload.soe.ucsc.edu/gbdb/hs1/bbi/cpgIslandExt.bb) 15.

Hg38ToHs1.over.chain from UCSC (https://hgdownload.soe.ucsc.edu/goldenPath/hg38/liftOver/hg38ToHs1.over.chain.gz) 16. The Relome browser containing the two Relome filter results and the top 5 utilized NTV3 sequence to function features per position is available at: https://relome.org/.

## Acknowledgements

This project was granted access to the JUPITER supercomputer through the JUPITER Research and Early Access Program (JUREAP) and the Gauss Centre for Supercomputing e.V. (GCS). JUPITER is funded by the EuroHPC Joint Undertaking, the German Federal Ministry of Research, Technology, and Space, and the Ministry of Culture and Science of North Rhine-Westphalia. Access to JUWELS and JURECA was granted by GCS and partly via the John von Neumann Institute for Computing. JUWELS is supported by GCS through funding by the German Federal Ministry of Research, Technology and Space and the Ministry of Culture and Science of the State of North Rhine-Westphalia. J.K. is funded by the Clinician Scientist program of the Faculty of Medicine, Uniklinik RWTH-Aachen.

## Author contributions

J.K. and I.K. conceived the study. J.K. performed computational workflows on JUPITER. M.Bod. supervised and coordinated high-performance computing activities. J.K. performed the genome-wide *in silico* perturbation analyses, and J.K. and T.I. generated the nucleotide-dependency maps. J.K., M.D. and T.I. performed downstream analyses including graph preprocessing and graph-based machine learning analyses. M.D. and J.K. developed the Relon detection methodology and performed statistical analyses. M.D. designed and implemented the Relome browser. F.K. performed splicing analyses and contributed to data evaluation. M.Beg. analysed non-coding and splice-related variants. M.E. led the clinical genetics assessment and interpretation of disease-associated genomic findings. Z.Z. prepared datasets for downstream modelling. J.K. and I.K. supervised the project. I.K., M.D. and J.K. wrote, edited and revised the manuscript with input from all authors. All authors read and approved the final manuscript.

## Funding

J.K. is supported by the Clinican Scientist Program of the Uniklinik RWTH Aachen.

## Competing interests

M.D. is also employed by scieneers GmbH, which was not involved in or financially supported this study. The remaining authors declare no competing interests.

## Supplementary information

### Supplementary Figures

**Supplementary Fig. 1.**
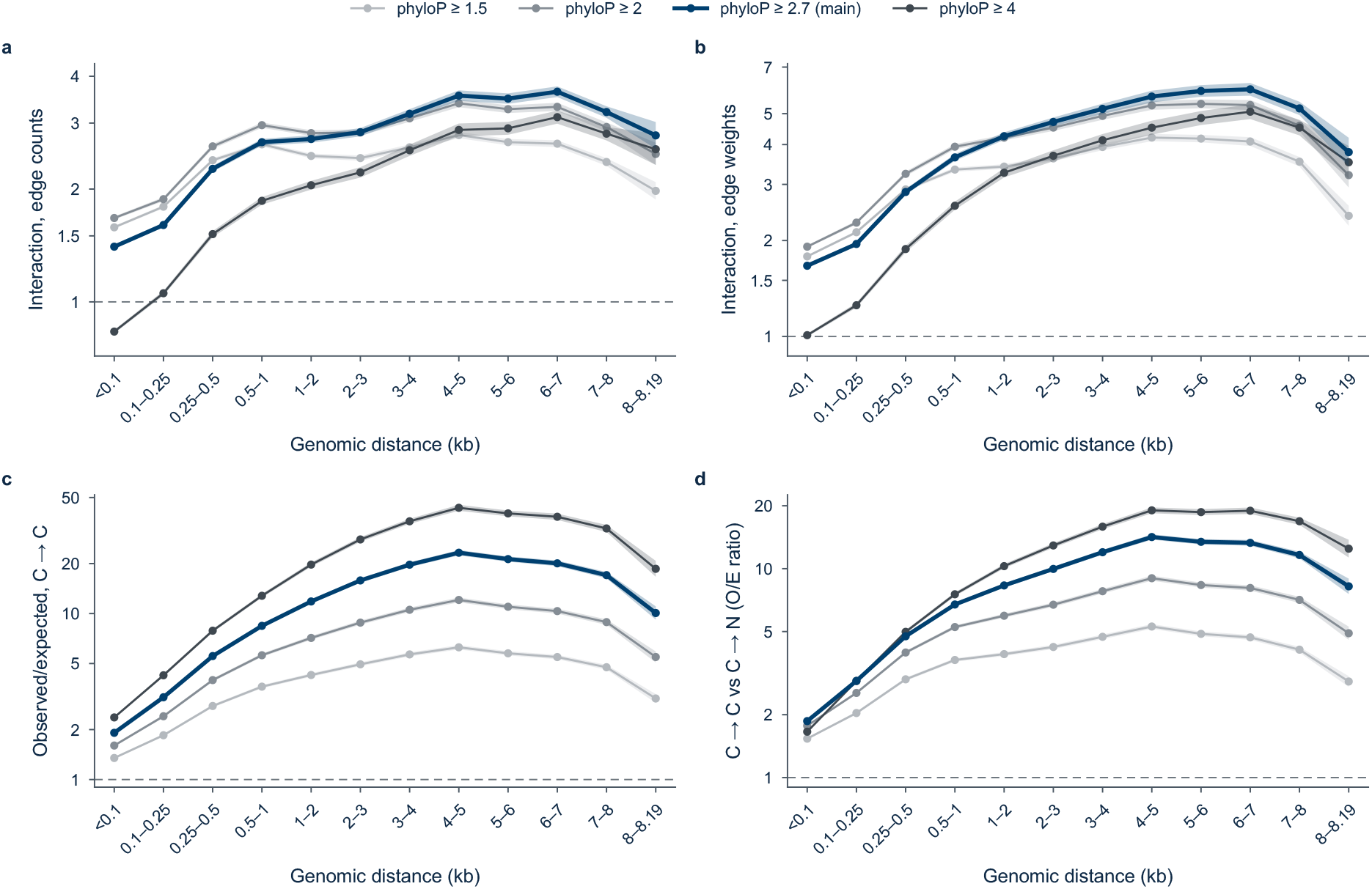
Conservation-associated nucleotide dependencies are robust across phyloP thresholds. Sensitivity analysis of conservation-associated dependency enrichment using four phyloP thresholds (≥1.5, ≥2, ≥2.7 and ≥4). Positions were classified as conserved (C) when their phyloP score met or exceeded the respective threshold, and as non-conserved (N) otherwise. Observed-to-expected (O/E) ratios measure dependency enrichment relative to expectations based on candidate position pairs matched for genomic distance within 500-kb blocks. **a,b**, Conservation-specific interaction across genomic distances for edge counts (a) and summed edge weights (b), calculated as (O/E C→C × O/E N→N) / (O/E C→N × O/E N→C). This interaction measures the association between source and target conservation beyond their independent effects on dependency frequency or strength; values above one indicate positive interaction. **c**, O/E ratios for edge counts between two conserved positions (C→C). **d**, Ratio of C→C to C→N enrichment, calculated as O/E C→C divided by O/E C→N, comparing the relative enrichment of these two dependency types. The principal patterns observed at the primary threshold (phyloP ≥2.7) were reproduced across alternative thresholds (a,b). For C→C enrichment and its comparison with C→N dependencies (c,d), enrichment generally increased with more stringent conservation thresholds.

**Supplementary Fig. 2.**
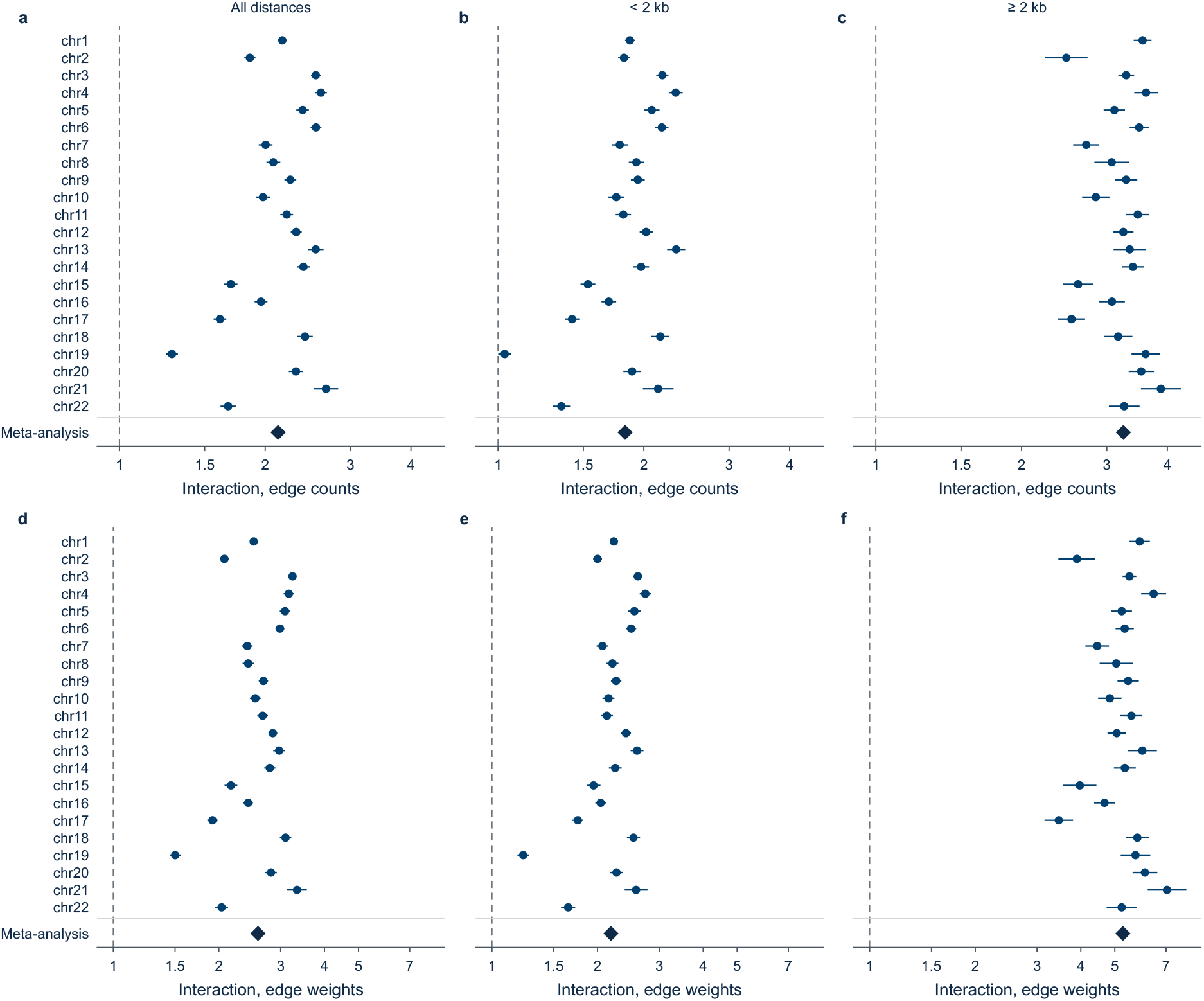
Conservation-specific nucleotide dependencies are observed across all autosomes. Chromosome-specific estimates of the interaction between source and target conservation, shown for all 22 human autosomes and their combined meta-analysis. Positions were classified as conserved (C; phyloP ≥2.7) or non-conserved (N). Observed-to-expected (O/E) ratios quantify dependency enrichment relative to expectations based on candidate position pairs matched for genomic distance within 500-kb blocks. Conservation-specific interaction was calculated as (O/E C→C × O/E N→N) / (O/E C→N × O/E N→C), accounting for the independent associations of source and target conservation with dependency frequency or strength. Values above one indicate preferential coupling between conserved positions beyond these individual effects. **a–c**, Interaction estimates based on edge counts across all evaluated genomic distances (a), distances below 2 kb (b) and distances of at least 2 kb (c). **d–f**, Corresponding interaction estimates based on summed edge weights across all distances (d), below 2 kb (e) and at least 2 kb (f). Each chromosome-specific estimate combines the respective genomic distance bins. Points indicate interaction estimates; horizontal lines show 95% confidence intervals and diamonds represent the combined meta-analysis across autosomes. Dashed vertical lines indicate an interaction value of one (no interaction). Conservation-specific interactions were consistently positive across autosomes, with stronger effects at genomic distances of at least 2 kb.

**Supplementary Fig. 3.**
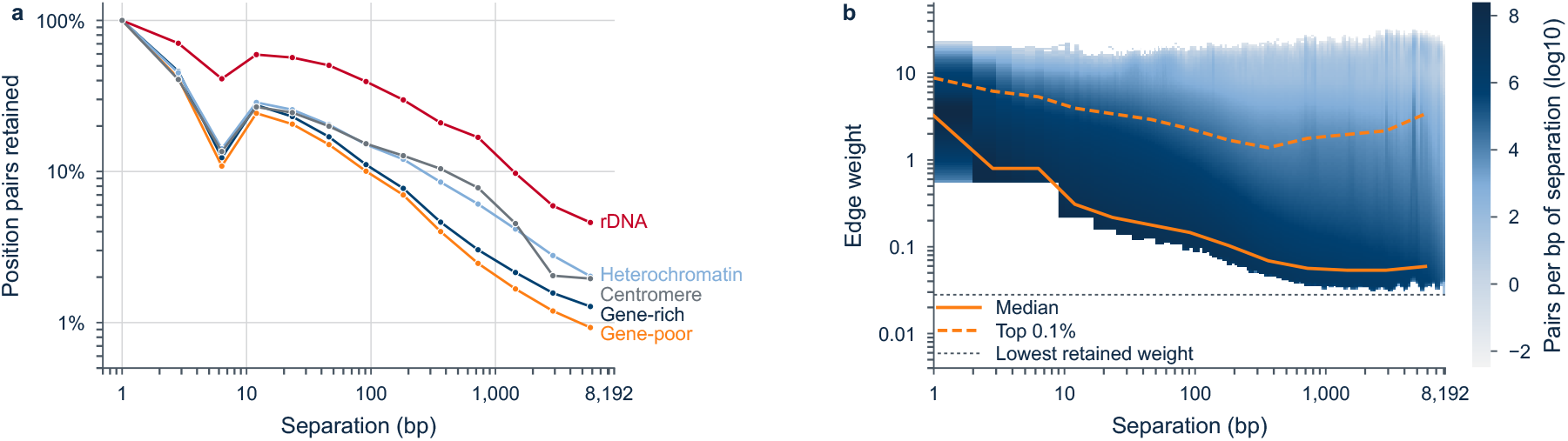
Distance-dependent nucleotide dependencies after exclusion of repetitive sequences. Analysis of nucleotide dependencies after excluding position pairs in which either nucleotide overlaps an annotated RepeatMasker element or a non-B DNA motif (sequences capable of adopting alternative DNA conformations). Annotated RNA genes were retained without masking. Analyses correspond to Fig. 3a,b but are restricted to the remaining, unmasked position pairs. **a**, Dependency retention across genomic distances and chromosomal contexts, defined as the proportion of eligible nucleotide pairs for which a dependency edge was retained after filtering. Chromosomal contexts include gene-rich, gene-poor, heterochromatic, centromeric and ribosomal DNA (rDNA) regions. Following masking, eligible pairs represented 23% of the original candidate pairs in gene-rich and gene-poor regions, 27% in rDNA, 3.5% in heterochromatic regions and 0.05% in centromeric regions (4.1 × 10⁸ pairs). Estimates for centromeric regions should therefore be interpreted cautiously, as they are derived from relatively few remaining genomic loci. **b**, Distribution of retained dependency weights across genomic separations. Solid orange lines indicate median edge weights, and dashed orange lines indicate the strongest 0.1% of weights at each separation. The dotted horizontal line marks the minimum retained edge weight of 0.028. Blue shading represents the number of nucleotide pairs contributing to each combination of genomic separation and edge weight, expressed on a log₁₀ scale. Each nucleotide pair was counted only once, irrespective of its occurrence in overlapping sequence windows; consequently, the colour scale is not directly comparable with Fig. 3b. Both panels use logarithmic axes.

**Supplementary Fig. 4.**
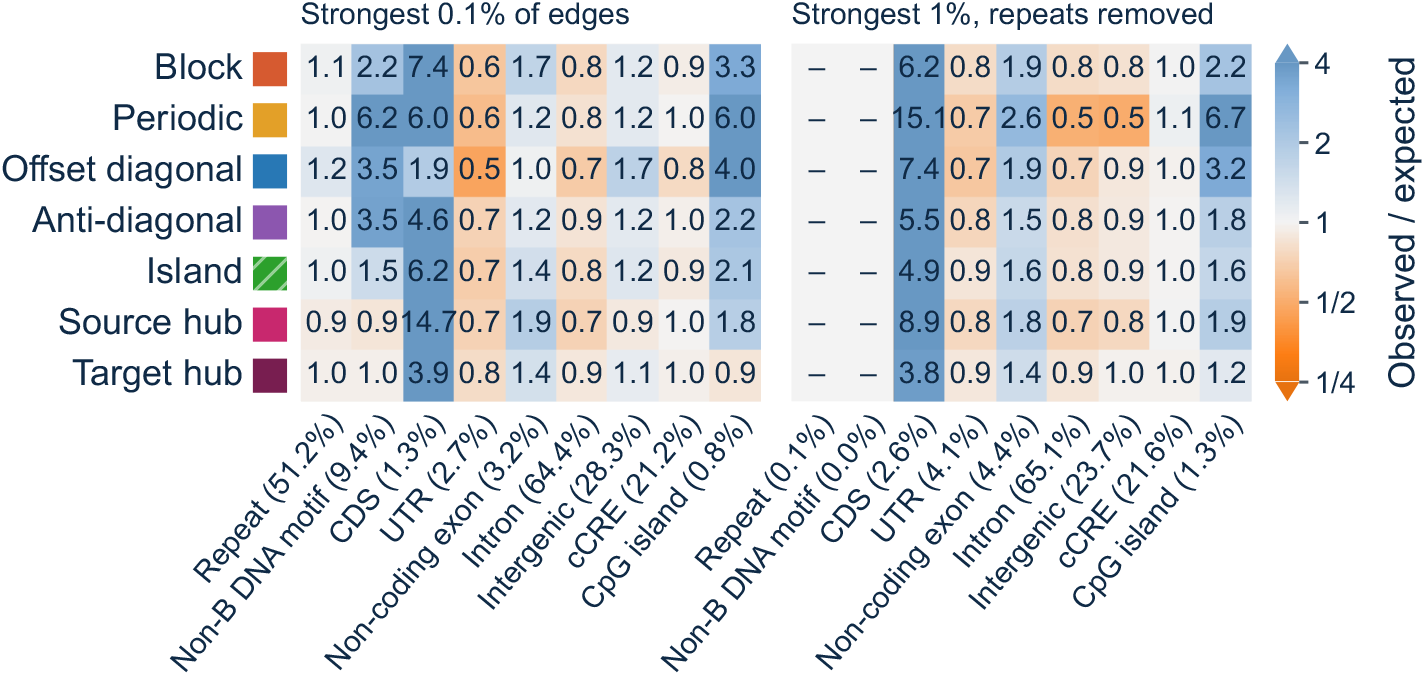
Relon candidates show distinct associations with genomic elements. Enrichment of genomic annotations across the seven Relon classes, calculated for individual Relon candidates before merging overlapping structures. Analyses were restricted to euchromatic regions. Each cell shows the observed-to-expected (O/E) enrichment of Relon-candidate bases overlapping a given genomic annotation, relative to the genomic coverage of that annotation in the corresponding analysis background. Values above one indicate enrichment, whereas values below one indicate depletion. **Left**, Relon candidates identified using the strongest 0.1% of dependency edges, retaining repeat-associated dependencies. **Right**, Relon candidates identified using the strongest 1% of edges after exclusion of repeat- and non-B-DNA-associated dependencies, considering only unmasked positions. Genomic annotations include repetitive elements (RepeatMasker), non-B DNA motifs (sequences capable of adopting alternative DNA conformations), coding sequences (CDS), untranslated regions (UTRs), non-coding exons, introns, intergenic regions, candidate cis-regulatory elements (cCREs) and CpG islands. Percentages in parentheses indicate the genomic coverage of each annotation within the corresponding analysis background. Annotations covering less than 0.5% of the analysed sequence were not tested (dashes). Colours indicate log₂-transformed O/E ratios, with the colour scale capped at fourfold enrichment or depletion; numerical values show the corresponding O/E ratios. Analyses were performed across all 24 chromosomes of the T2T-CHM13 reference genome.

**Supplementary Fig. 5.**
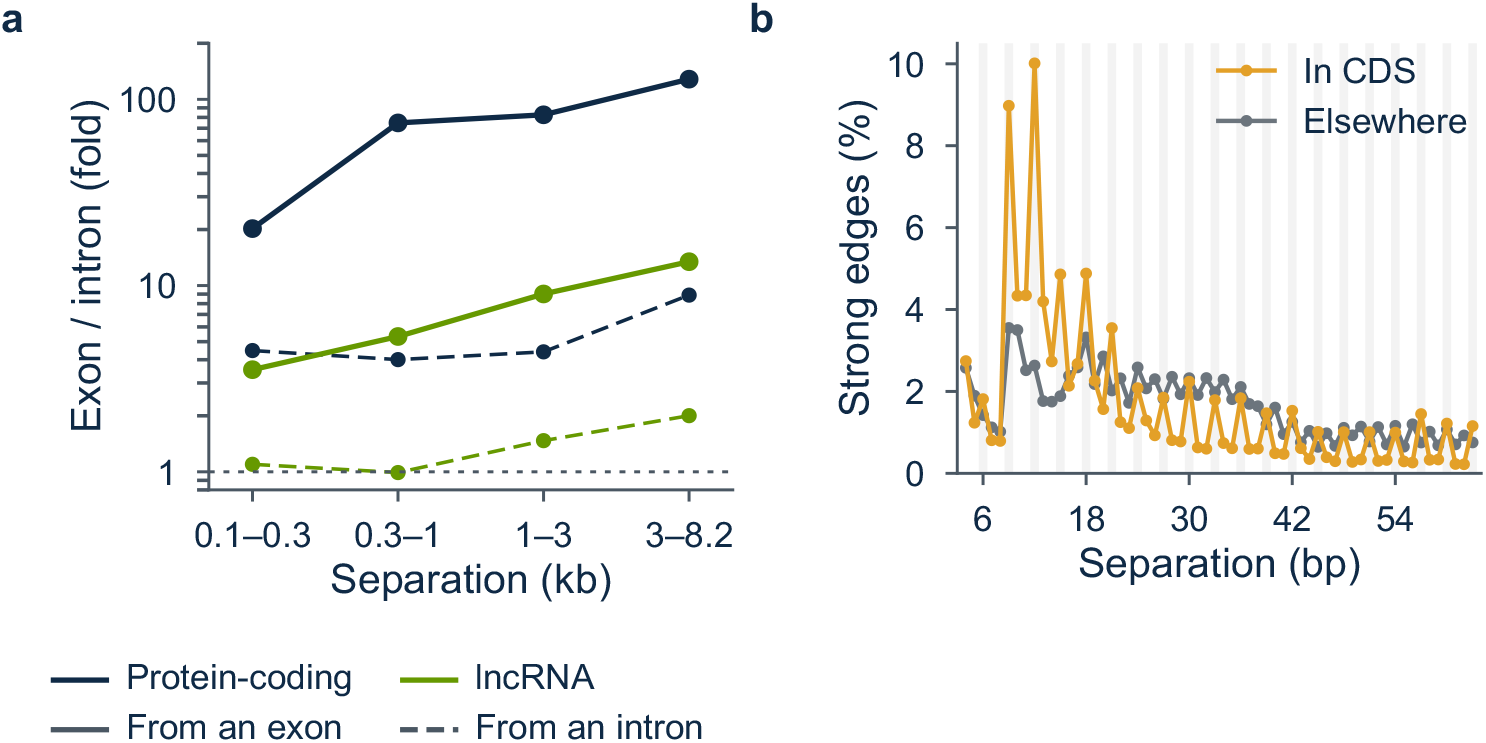
Exon connectivity and three-base periodicity with repeats retained. Analysis of gene-associated nucleotide dependencies under the strongest 0.1% edge filter, retaining repeat-associated dependencies. **a**, Enrichment of exon-to-exon dependencies relative to exon-to-intron dependencies across four genomic distance ranges, shown separately for protein-coding genes (black) and long non-coding RNA (lncRNA) genes (green). Enrichment is calculated as the ratio of dependency frequencies, accounting for the number of possible nucleotide pairs in each category. Solid lines indicate exon-to-exon enrichment, whereas dashed lines show the corresponding intron-based controls. Values above one indicate preferential connectivity. Exon-to-exon enrichment in protein-coding genes reached approximately 128-fold at distances of 3–8.2 kb. Analyses used one representative transcript per multi-exon gene: MANE Select transcripts for 17,073 protein-coding genes and the transcripts containing the greatest number of exons for 24,723 lncRNA genes. Only dependencies with both endpoints within the same Relon were included. **b**, Distribution of genomic separations (4–63 bp) among strong dependency edges in periodic Relon candidates predominantly overlapping coding sequence (CDS; orange, at least 50% of candidate bases) compared with all other periodic candidates (grey). The y-axis indicates the percentage of strong edges occurring at each separation. Separations divisible by three are highlighted, revealing the characteristic three-base periodicity associated with coding sequence.

**Supplementary Fig. 6.**
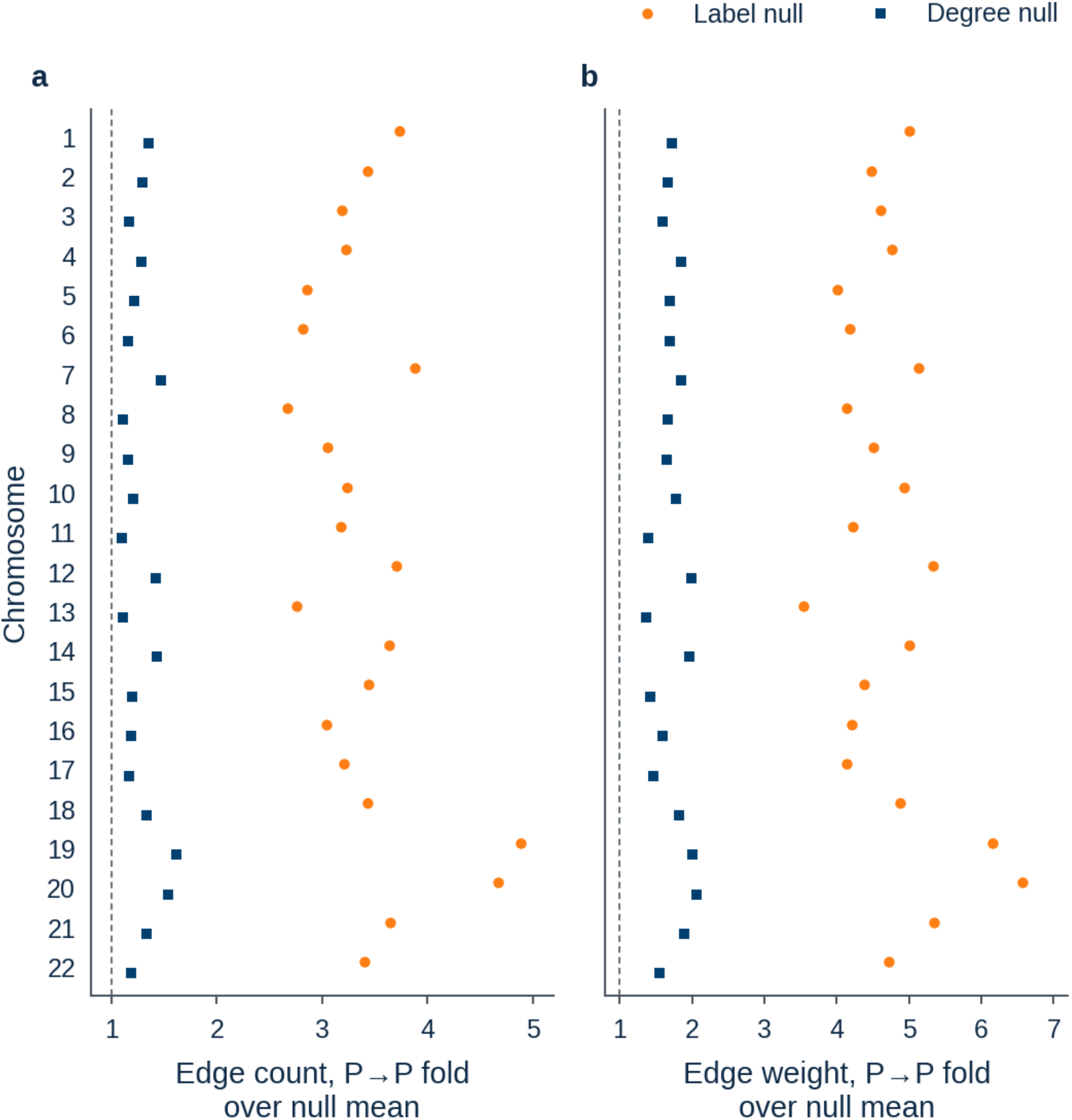
Dependencies between pathogenic variants exceed permutation-based expectations across all autosomes. Enrichment of nucleotide dependencies connecting positions harbouring pathogenic (P) ClinVar variants, evaluated separately for each of the 22 human autosomes. Enrichment was calculated as the observed fraction of pathogenic-to-pathogenic (P→P) dependencies divided by the mean fraction obtained from 5,000 permutations. Two null models were compared. In the label-permutation null (orange), pathogenic and benign variant labels were randomly shuffled across genomic positions while preserving the dependency network. In the degree-matched null (blue), labels were shuffled only among positions with similar numbers of outgoing and incoming dependencies (out-degree and in-degree, respectively), thereby accounting for differences in overall nucleotide connectivity. **a**, Enrichment of P→P edge counts relative to the two null models. **b**, Corresponding enrichment based on summed edge weights, which quantify dependency strength. Each point represents one autosome. Dashed vertical lines indicate a ratio of one, corresponding to no enrichment relative to the respective null model. P→P dependencies exceeded both null expectations across all autosomes, with substantially reduced enrichment after accounting for nucleotide connectivity.

**Supplementary Fig. 7.**
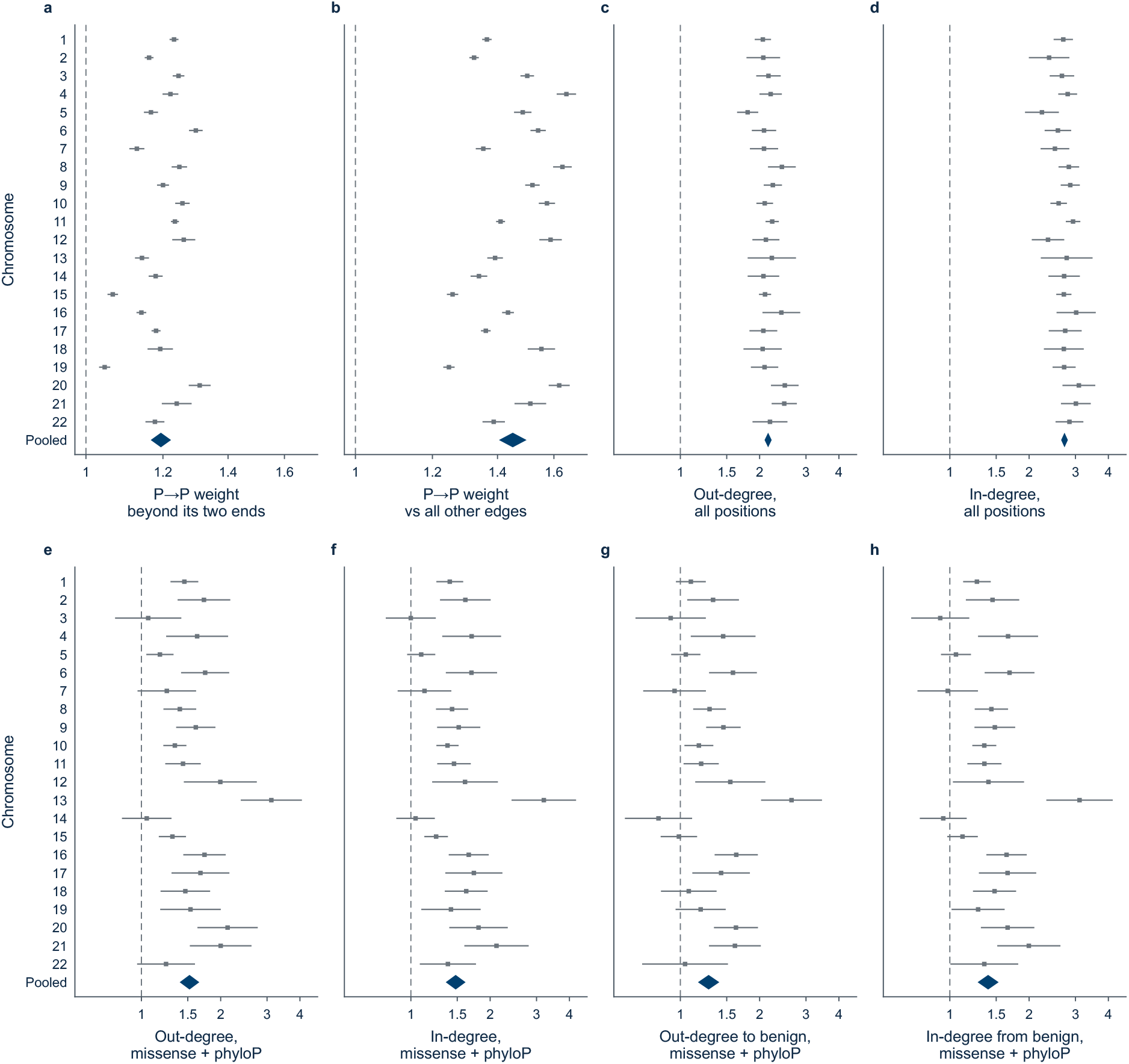
Pathogenic variants show increased nucleotide connectivity and dependency strength across autosomes. Chromosome-specific and pooled analyses of nucleotide dependencies involving positions harbouring pathogenic (P) or benign (B) ClinVar variants. Positions with conflicting clinical classifications were excluded. Multiple edges connecting the same directed pair of positions were merged, retaining their mean weight. Grey squares indicate estimates for each of the 22 autosomes, with horizontal lines showing 95% confidence intervals (CIs). Blue diamonds represent pooled estimates with 95% CIs, calculated using a DerSimonian–Laird random-effects meta-analysis of log-transformed estimates. This approach combines chromosome-specific estimates while accounting for variation between chromosomes. All x-axes use logarithmic scales; dashed vertical lines indicate a ratio of one (no enrichment). **a**, Enrichment of dependency weights between two pathogenic positions (P→P) beyond the independent effects of pathogenicity at the source and target positions (interaction term): 1.19-fold (95% CI, 1.17–1.22; I² = 98%). **b**, Enrichment of P→P dependency weights relative to all other dependency types, without adjustment for the independent effects of source and target pathogenicity: 1.45-fold (95% CI, 1.40–1.50; I² = 99%). **c,d**, Relative connectivity of pathogenic versus benign positions across all eligible ClinVar positions, measured by out-degree (c; 2.15-fold, 95% CI, 2.09– 2.22; I² = 48%) and in-degree (d; 2.73-fold, 95% CI, 2.65–2.81; I² = 36%). Out-degree denotes the number of dependencies originating from a position, whereas in-degree denotes the number of dependencies directed towards it. **e,f**, Corresponding out-degree (e; 1.52-fold, 95% CI, 1.40–1.66; I² = 78%) and in-degree (f; 1.48-fold, 95% CI, 1.36–1.61; I² = 80%) comparisons restricted to missense variant positions and adjusted for evolutionary conservation (phyloP) using a natural cubic spline to accommodate nonlinear associations. **g**, Out-degree comparison as in e, counting only dependencies directed towards benign positions (1.28-fold, 95% CI, 1.17–1.41; I² = 79%). **h**, In-degree comparison as in f, counting only dependencies originating from benign positions (1.40-fold, 95% CI, 1.28–1.53; I² = 81%). The I² statistic quantifies between-chromosome heterogeneity in the meta-analysis. Heterogeneity was particularly high for the P→P weight comparisons (a,b; I² = 98–99%), with 95% prediction intervals of 1.06–1.35 (a) and 1.23–1.71 (b).

**Supplementary Fig. 8.**
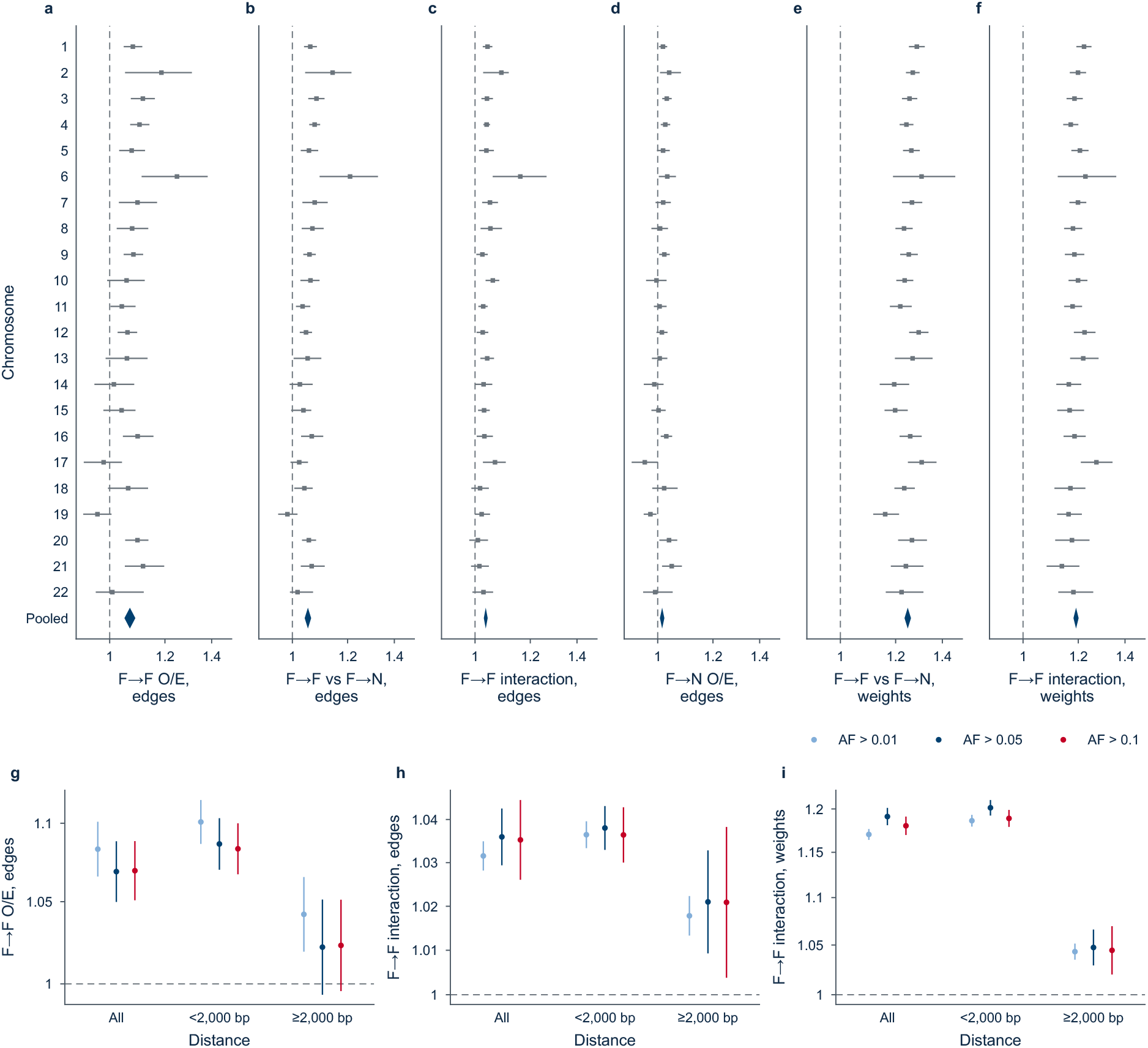
Frequent variants show distance-dependent enrichment in nucleotide dependencies. Analysis of nucleotide dependencies between genomic positions harbouring frequent variants in gnomAD, evaluated across the 22 human autosomes. Positions were classified as frequent (F) when their allele frequency (AF) exceeded the specified threshold (primarily AF >0.05), and as non-frequent (N) otherwise. Observed-to-expected (O/E) ratios quantify enrichment relative to the number or summed weight of dependencies expected from the proportion of candidate source–target pairs in each category, matched for genomic distance within 500-kb blocks. This accounts for the genomic clustering of frequent variants. Dependency direction distinguishes source positions, where nucleotides were computationally substituted, from target positions, where the resulting model responses were measured. **a**, O/E enrichment of edge counts between frequent positions (F→F). **b**, Relative enrichment of F→F compared with F→N edge counts, calculated as O/E(F→F) / O/E(F→N), indicating whether frequent source positions preferentially connect to frequent rather than non-frequent targets. **c**, Conservation-independent interaction between frequent source and target positions, calculated for edge counts as [O/E(F→F) × O/E(N→N)] / [O/E(F→N) × O/E(N→F)]. This interaction accounts for the independent associations of source and target frequency status with dependency formation. **d**, O/E enrichment of edge counts from frequent to non-frequent positions (F→N). **e,f**, Corresponding F→F versus F→N enrichment (e) and source–target interaction (f), calculated using summed dependency weights rather than edge counts. **a–f**, Grey squares show chromosome-specific estimates with 95% confidence intervals (CIs); blue diamonds indicate pooled estimates across all 22 autosomes, using the primary threshold of AF >0.05 and all evaluated genomic distances. **g–i**, Pooled estimates with 95% CIs across alternative frequency thresholds (AF >0.01, >0.05 and >0.1), shown for all distances combined and separately for distances below 2 kb and at least 2 kb. Panels show F→F O/E enrichment for edge counts (g), F→F interaction for edge counts (h) and F→F interaction for summed edge weights (i). Dashed lines indicate a ratio of one, corresponding to no enrichment or interaction. Frequent-variant positions showed modest enrichment in dependency counts, whereas enrichment in dependency weights was more pronounced, particularly at genomic distances below 2 kb.

**Supplementary Figure 9:**
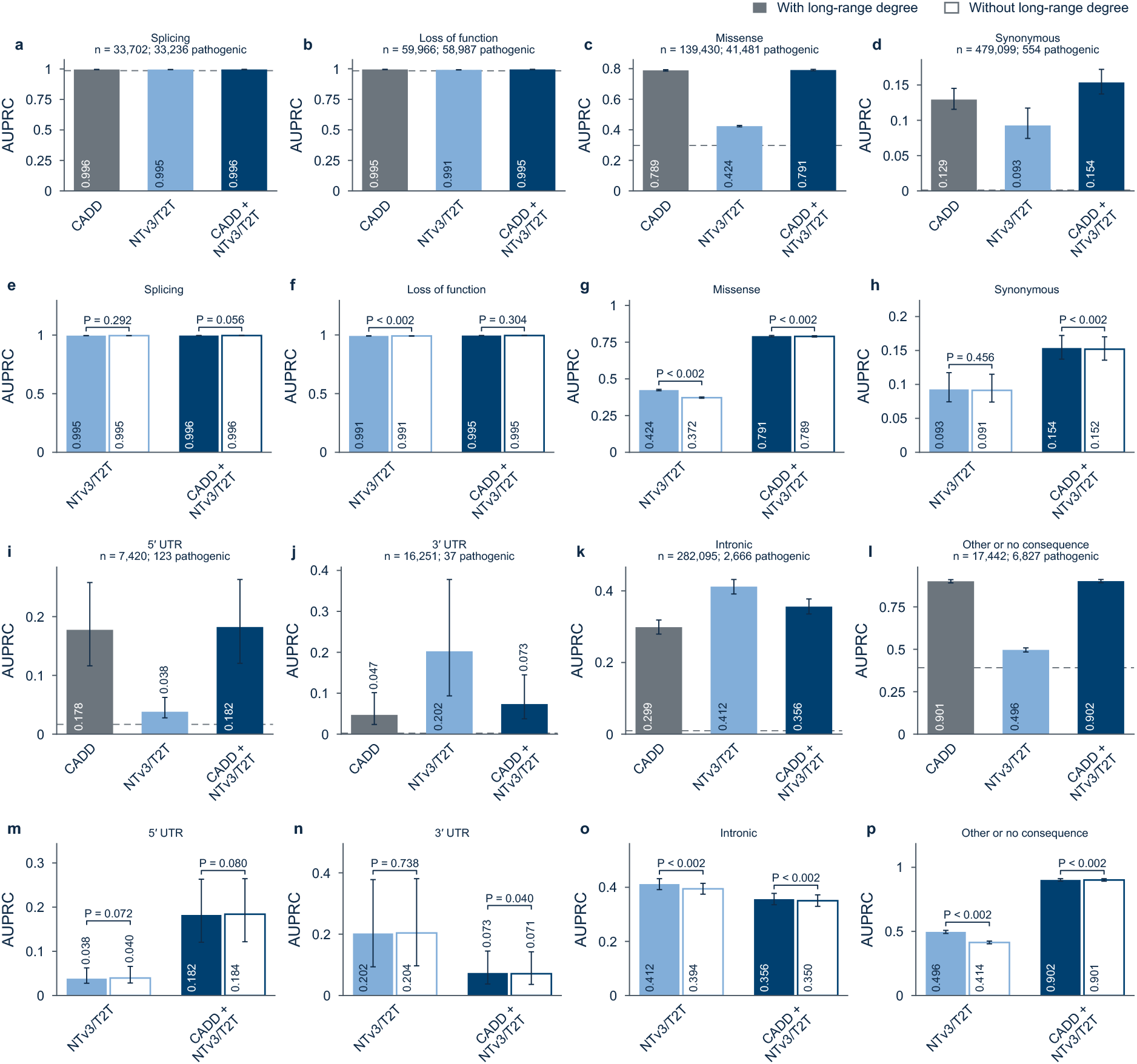
Combining NTv3/T2T features with CADD improves classification of noncoding and synonymous ClinVar variants. AUPRC of CADD, NTv3/T2T and CADD + NTv3/T2T per consequence class (**a–d**, **i–l**), and of NTv3/T2T and CADD + NTv3/T2T with (filled) and without (open) long-range degree (**e–h**, **m–p**). Classes: splice site (a, e), loss of function (b, f), missense (c, g), synonymous (d, h), 5′ UTR (i, m), 3′ UTR (j, n), intronic (k, o), other (l, p). Error bars, 95% CIs from 1,000 bootstrap resamples; P values, paired bootstrap test; dashed line, fraction of pathogenic variants. In this evaluation variant consequence classes were treated as exclusive, following the priority defined in the methods section.

**Supplementary Fig. 10.**
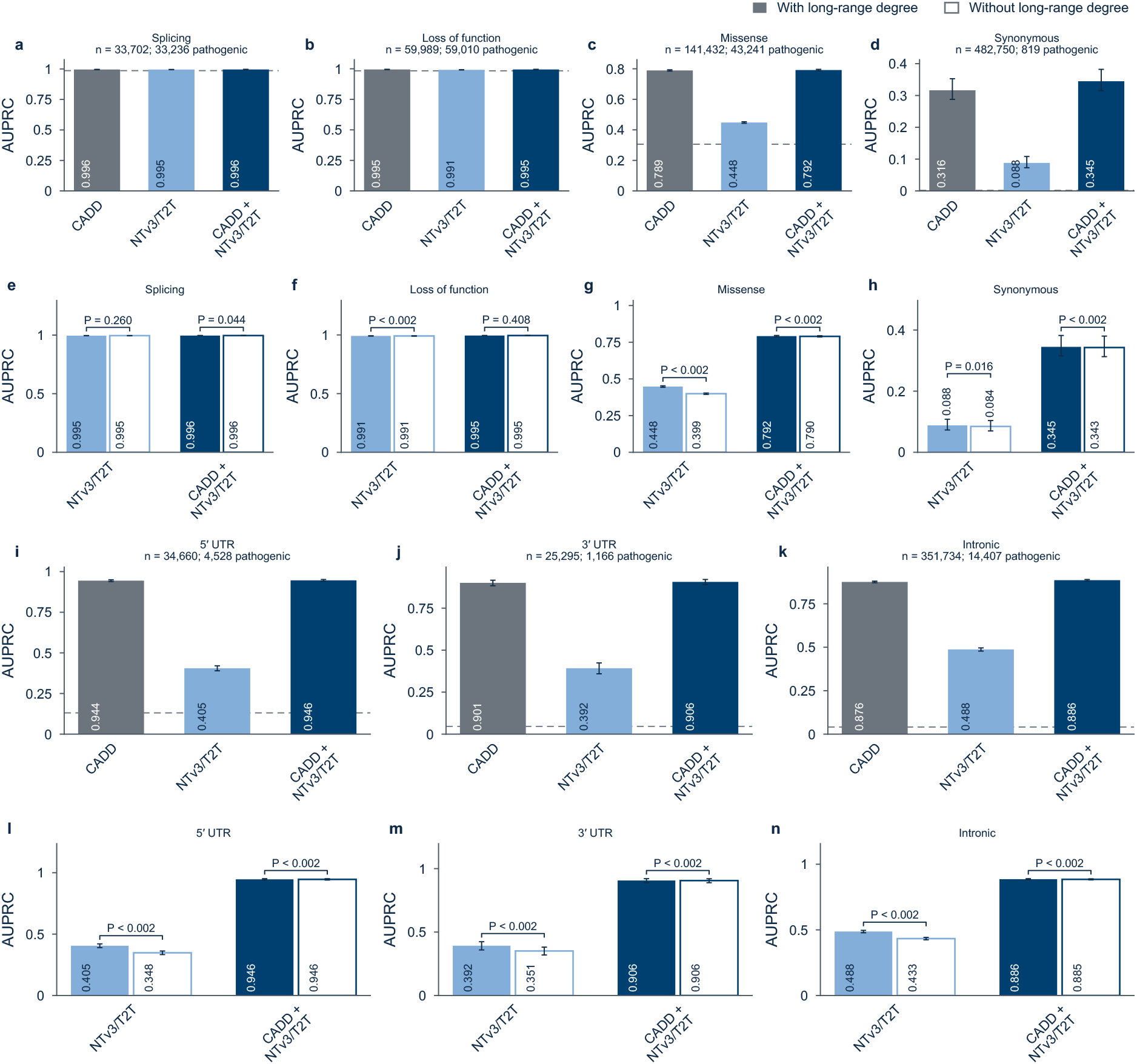
: Combining NTv3/T2T features with CADD improves classification of noncoding and synonymous ClinVar variants. AUPRC of CADD, NTv3/T2T and CADD + NTv3/T2T per consequence class (**a–d**, **i–l**), and of NTv3/T2T and CADD + NTv3/T2T with (filled) and without (open) long-range degree (**e–h**, **m–p**). Classes: splice site (a, e), loss of function (b, f), missense (c, g), synonymous (d, h), 5′ UTR (i, m), 3′ UTR (j, n), intronic (k, o), other (l, p). Error bars, 95% CIs from 1,000 bootstrap resamples; P values, paired bootstrap test; dashed line, fraction of pathogenic variants. In this evaluation variant classes were treated as non-exclusive, therefore one variant could have belonged to multiple variant consequence classes.

**Supplementary Fig. 11.**
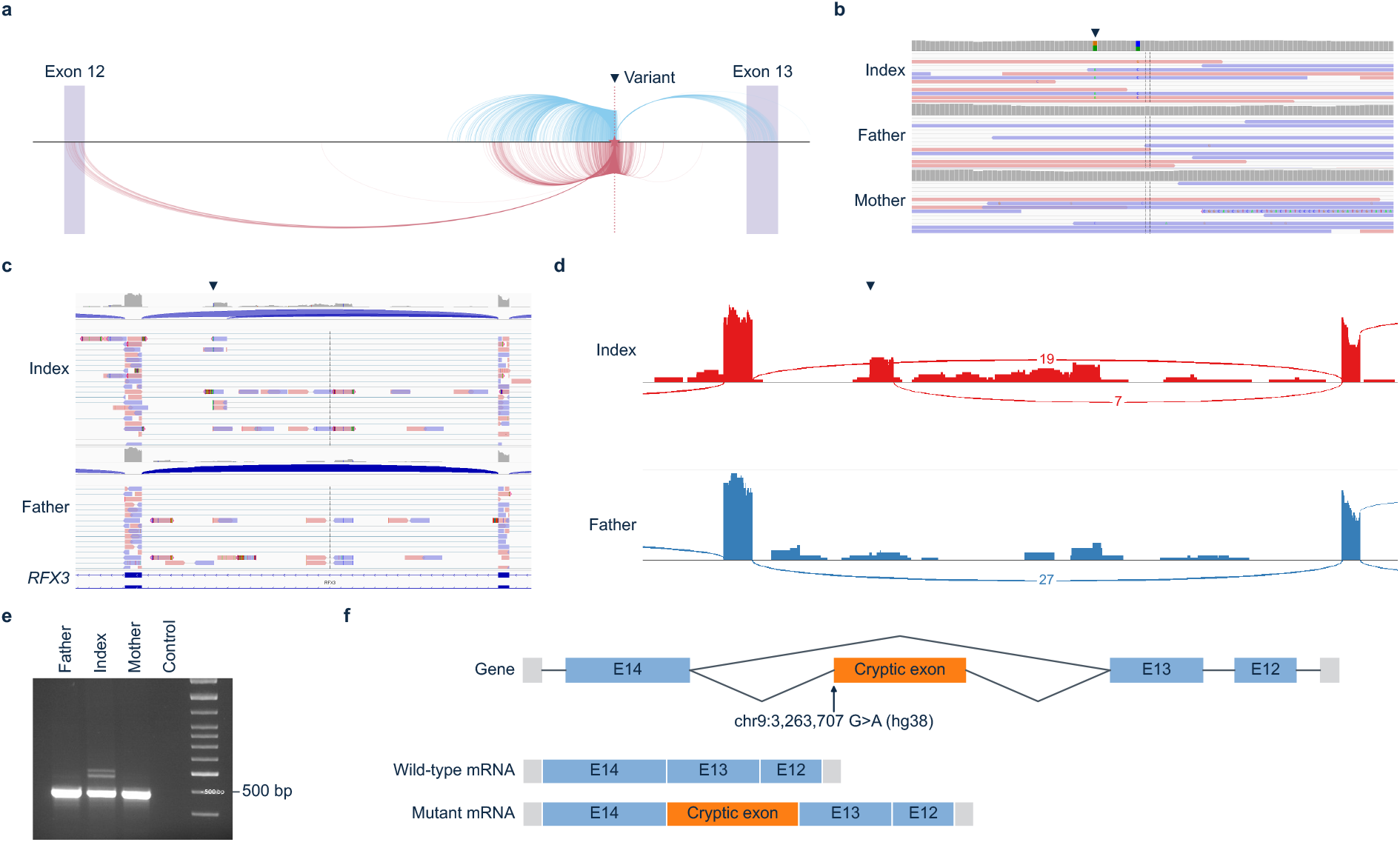
Nucleotide dependencies and aberrant splicing at the RFX3 locus. **a**, Nucleotide-dependency landscape surrounding a deep-intronic RFX3 variant (chr9:3,267,537; T2T-CHM13v2.0). Directed dependency arcs connect the intronic region harbouring the variant with neighbouring sequences, including flanking exons. Vertical shaded regions indicate annotated exons, and the dashed line marks the variant position. **b**, Genomic sequencing read alignments of the affected individual (index, top) and both parents (middle (father), bottom, mother), visualized using the Integrative Genomics Viewer (IGV), showing the intronic de novo variant in RFX3. **c**, RNA-sequencing read alignments across the affected RFX3 intron, illustrating intronic read coverage and splice-junction patterns. **d**, Corresponding sashimi plots showing RNA-sequencing coverage and splice junctions across the affected region. The upper (red) track displays additional splice junctions involving intronic sequence, whereas the lower (blue) track predominantly shows the canonical splice junction connecting the flanking exons. Coverage profiles indicate mapped RNA-sequencing reads, and arcs represent splice junctions supported by split reads. **e**, Reverse-transcription PCR (RT-PCR) analysis of RNA isolated from peripheral blood of the affected individual (middle lane) and the parental samples (adjacent, left, right) and a no template control (NTC). RNA was converted into complementary DNA (cDNA), followed by PCR amplification using primers in the flanking exons. Agarose gel electrophoresis shows a shared product corresponding to the normally spliced transcript in all samples, together with additional higher-molecular-weight products in the affected individual (middle lane). A DNA size marker is shown on the right. **f**, Schematic representation of RFX3 splicing, illustrating the normally spliced transcript and an aberrant transcript containing an additional 125-nucleotide cryptic exon. The deep-intronic variant is indicated at chr9:3,263,707G>A (GRCh38). Together, these findings demonstrate aberrant splicing associated with the intronic RFX3 variant, while the nucleotide-dependency landscape independently reveals model-derived relationships between the affected intronic region and neighbouring exons.

### Supplementary Tables

**Supplementary Table 1.** Rules for Relon classification. Seven Relon classes defined by the geometry and directionality of nucleotide dependencies. Candidates were assigned to the first matching class in the order shown. Widths denote source and target intervals; correlations describe their positional correspondence. Further details are provided in Methods.

| Order | Class | Pattern |
| --- | --- | --- |
| 1 | Source hub | The source stretch is at most a quarter as wide as the target stretch: a narrow region whose perturbation affects a much broader one |
| 2 | Target hub | The converse: predictions in a narrow region depend on perturbations across a much broader one |
| 3 | Anti-diagonal | Source and target positions run in opposite directions (correlation $\leq -0.5$), as in a mirror image; the two stretches may meet at the diagonal |
| 4 | Offset diagonal | Two separate stretches whose positions run in the same direction (correlation $\geq 0.5$), like two aligned copies |
| 5 | Island | Two separate stretches without consistent order between their positions |
| 6 | Periodic | At least half of the edges fall on the three most frequent separations: dependencies recur at fixed intervals |
| 7 | Block | All remaining candidates: a dense patch on the diagonal |

**Supplementary Table 2.** Representative examples of the seven Relon classes. Genomic coordinates and identifiers of Relons shown in Fig. 4, selected from chromosome 21 of the T2T-CHM13v2.0 reference genome using the strongest 0.1% edge filter. Links open the corresponding regions in the Relome browser.

| Class | Relon | Region (chr21) | Relome browser |
| --- | --- | --- | --- |
| Block | R17546 | 45,077,348–45,079,790 | https://relome.org/?locus=chr21:45077348-45079790&tracks=relons&shapes=block |
| Periodic | R17147 | 44,408,159–44,409,122 | https://relome.org/?locus=chr21:44408159-44409122&tracks=relons&shapes=periodic |
| Offset diagonal | R2742 | 11,769,564–11,772,996 | https://relome.org/?locus=chr21:11769564-11772996&tracks=relons&shapes=offset+diagonal |
| Anti-diagonal | R16454 | 43,108,585–43,111,072 | https://relome.org/?locus=chr21:43108585-43111072&tracks=relons&shapes=anti-diagonal |
| Island | R7569 | 23,600,793–23,601,524 | https://relome.org/?locus=chr21:23600793-23601524&tracks=relons&shapes=island |
| Source hub | R15360 | 41,102,085–41,102,465 | https://relome.org/?locus=chr21:41102085-41102465&tracks=relons&shapes=source+hub |
| Target hub | R12682 | 34,990,264–34,992,631 | https://relome.org/?locus=chr21:34990264-34992631&tracks=relons&shapes=target+hub |

### Supplementary Notes

#### Supplementary Note 1 | Relon detection and classification

##### Dependency maps and background expectations

Each 8,192-bp genomic sequence window was represented as a nucleotide-dependency matrix, with perturbed source positions along the rows and responding target positions along the columns. Strong dependencies were identified separately by genomic separation and cytoband class to account for differences in dependency frequency across distances and chromosomal environments. For each source–target position pair, we calculated the expected frequency of strong dependencies from pairs with the same separation and cytoband class. Relon detection was based on regions containing more strong dependencies than expected under this background model.

##### Identification of Relon candidates

Dependency matrices were scanned using square boxes of 4, 16, 64, 128 and 256 bp to identify enriched structures at different spatial scales. A box qualified as a seed if it contained at least three strong edges and showed at least eightfold enrichment for 4-bp boxes or fourfold enrichment for larger boxes. Candidates were extended into neighbouring boxes containing dependencies, provided their enrichment remained at least twofold. To distinguish recurrent structures from random edge clusters, we generated simulated dependency maps preserving the expected edge frequencies by genomic separation and cytoband class. Candidates were retained only if their most enriched seed exceeded the threshold reached by one in ten million random boxes of the corresponding size and cytoband class. Each retained candidate was then trimmed to the source and target positions carrying its edges, defining its genomic intervals.

##### Classification and merging

Candidates were assigned to seven Relon classes using a hierarchical set of geometric rules (Supplementary Table 1). The classification distinguishes patterns of directional connectivity, including source and target hubs, anti-diagonals, offset diagonals, islands, periodic structures and blocks. Rules were applied sequentially, and each candidate was assigned to the first matching class. Because dependencies are directed, relationships from position A to B and from B to A were evaluated independently.

Individual dependency architectures could generate multiple overlapping candidates, for example at different box sizes, in opposite edge directions or as partially overlapping fragments. We therefore removed candidates representing primarily short-range diagonal thickening (predominantly 2–3-bp separations), replaced broader candidates with nested candidates retaining at least 80% of their edges, and merged candidates whose genomic intervals overlapped or were connected by a strong dependency. Each resulting Relon inherited the class of its constituent candidate containing the greatest number of edges. Representative examples of all seven Relon classes, together with their genomic coordinates and direct links to the Relome browser, are provided in Supplementary Table 2.

#### Supplementary Note 2 | Effects of alternative edge filters on Relon composition

Relon class composition differed between the strongest 0.1% edge filter, which retained repeat-associated dependencies, and the alternative filter selecting the strongest 1% of edges after excluding repeat- and non-B-DNA-associated dependencies.

Under the strongest 0.1% filter, at least half of the bases were located in regions excluded by the alternative filter for 86% of periodic Relons, 83% of anti-diagonals and 76% of offset diagonals, compared with 51–67% of Relons in the remaining classes. Approximately 90% of the predominantly masked periodic Relons and anti-diagonals overlapped mainly non-B DNA motifs rather than RepeatMasker elements.

Blocks showed a different pattern. Excluding predominantly masked Relons from the original 0.1% dataset would increase their relative abundance only from 15.8% to 17.8%, substantially less than the 32.6% observed under the alternative filter. Their increased representation therefore cannot be explained by repeat exclusion alone and is consistent with an additional contribution from the more permissive edge threshold. Moreover, at most 375,000 of the 1.21 million Relons identified under the alternative filter could have been carried over from the original analysis. Thus, differences between the two Relon landscapes reflect both sequence masking and changes in edge selection.

#### Supplementary Note 3 | Relons and genomic annotation

##### Class-specific genomic associations

To prevent large centromeric and heterochromatic Relons from dominating enrichment analyses, genomic annotation overlap was evaluated within euchromatin (Giemsa bands gneg and gpos). Offset diagonals were enriched in intergenic sequence (2.1-fold) but depleted in introns (0.5-fold), untranslated regions (UTRs; 0.4-fold) and candidate cis-regulatory elements (cCREs; 0.7-fold). Inverted repeats showed only modest enrichment in anti-diagonal Relons (2.0-fold). Periodic Relons were enriched in CpG islands under both edge filters (4.7- and 5.8-fold), while non-coding exons showed modest enrichment across most classes (up to 1.8-fold).

##### Effects of candidate merging

Class-specific genomic associations were also evaluated before merging overlapping Relon candidates (Supplementary Fig. 4). Because a merged Relon inherits the class of its largest candidate, merging can obscure associations detected for individual component candidates. In particular, source-hub candidates remained strongly enriched in coding sequence after repeat exclusion (8.9-fold), whereas the corresponding merged source hubs showed no enrichment (1.0-fold). The original analysis also identified differences between candidate-level and merged-level class attribution involving 21.2% and 33.9% of euchromatic Relon bases under the strongest 0.1% and repeat-excluding filters, respectively.

##### Relons beyond genomic annotation

Dark Relons were defined as Relons whose source and target positions showed no overlap with annotated genes (exons or introns), Repeat-Masker elements, cCREs, small open reading frames, non-B DNA motifs, Tandem Repeats Finder elements or CpG islands.

Under the strongest 0.1% edge filter, Dark Relons accounted for 1.7% of all Relons, ranging from 0.6% on chromosome 19 to 3.0% on chromosomes 13, 21 and Y. Islands and source or target hubs together accounted for 68% of Dark Relons.

To determine whether these architectures preferentially occurred in unannotated regions, each Relon was compared with a randomly shifted copy on the same chromosome within the same cytoband class. Dark Relons were less frequent than expected under this control (1.7% observed versus 4.9% expected). This depletion was particularly pronounced for periodic Relons (0.03-fold), anti-diagonals (0.16-fold) and blocks (0.22-fold), whereas islands and hubs were closer to random expectation (0.79–1.02-fold).

After repeat exclusion, this depletion was no longer observed: among Relons without masked bases, 17.6% were dark, compared with 17.0% of their randomly shifted controls. These analyses show that Relons are not generally enriched in unannotated regions, but that recurrent nucleotide-dependency architectures also occur extensively outside currently annotated genomic elements.

## Notes

https://relome.org/

### Competing Interest Statement

The authors have declared no competing interest.

### Summary of Updates

Chance of author list in the online version

